# AspWood: High-spatial-resolution transcriptome profiles reveal uncharacterized modularity of wood formation in *Populus tremula*

**DOI:** 10.1101/094060

**Authors:** David Sundell, Nathaniel R. Street, Manoj Kumar, Ewa J. Mellerowicz, Melis Kucukoglu, Christoffer Johnsson, Vikash Kumar, Chanaka Mannapperuma, Nicolas Delhomme, Ove Nilsson, Hannele Tuominen, Edouard Pesquet, Urs Fischer, Totte Niittylä, Bjöern Sundberg, Torgeir R. Hvidsten

**Affiliations:** Umeå Plant Science Center, Department of Plant Physiology, Umeå University, Umea, Sweden; Umeå Plant Science Centre, Department of Forest Genetics and Plant Physiology, Swedish University of Agricultural Sciences, Umeå, Sweden; Department of Ecology, Environment and Plant Sciences, Stockholm University, Sweden; Department of Chemistry, Biotechnology and Food Sciences, Norwegian University of Life Sciences, Ås, Norway

**Keywords:** Cambium growth, wood formation, transcriptomics, co-expression networks, paralogues, cell wall

## Abstract

Trees represent the largest terrestrial carbon sink and a renewable source of ligno-cellulose. There is significant scope for yield and quality improvement in these largely undomesticated species, and efforts to engineer new, elite varieties will benefit from an improved understanding of the transcriptional network underlying cambial growth and wood formation. We generated high-spatial-resolution RNA Sequencing data spanning the secondary phloem, vascular cambium and wood forming tissues. The transcriptome comprised 28,294 expressed, previously annotated genes, 78 novel protein-coding genes and 567 long intergenic non-coding RNAs. Most paralogs originating from the *Salicaceae* whole genome duplication were found to have diverged expression, with the notable exception of those with high expression during secondary cell wall deposition. Co-expression network analysis revealed that the regulation of the transcriptome underlying cambial growth and wood formation comprises numerous modules forming a continuum of active processes across the tissues. The high spatial resolution enabled identification of novel roles for characterised genes involved in xylan and cellulose biosynthesis, regulators of xylem vessel and fiber differentiation and lignification. The associated web resource (AspWood, http://aspwood.popgenie.org) integrates the data within a set of interactive tools for exploring the expression profiles and co-expression network.

## Introduction

Trees dominate forest ecosystems, with the majority of biomass residing in the wood of stems, branches and roots. Cambial growth (production of new secondary phloem and xylem cells by periclinal divisions) and wood formation (secondary xylem cell differentiation or xylogenesis) are initiated in the vascular cambium meristem (hereafter, cambium), which forms a cylindrical sheath of dividing cells within the stem (Barnett, 1981; Larson, 1994; Mellerowicz et al., 2001). Inwards, the cambium forms secondary xylem (wood) and outwards, secondary phloem cells are added to the growing stem. The cambium consists of stem cells (also referred to as initials) and their dividing derivatives (also referred to as xylem and phloem mother cells) (Johnsson and Fischer, 2016). The stem cells retain the capacity to divide over long periods of time (stem cell maintenance), whereas derivative cells divide over a few cell cycles. Before terminal differentiation into specialized cell types, all xylem and phloem cells undergo initial cell expansion and primary cell wall (PCW) biosynthesis. Derivatives at the outer cambial face differentiate into the different cell types forming the phloem, including sieve tube cells involved in long-range transport of photosynthates, ray and axial parenchyma cells, including companion cells, and phloem fibers with thickened and lignified cell walls (Evert and Eichhorn, 2006). In most angiosperm tree species, cambial derivatives on the xylem side differentiate into four major wood cell types; fibers that provide structural support, vessel elements for water and mineral transport, axial parenchyma cells for storage, and ray cells involved in radial transport and storage of photosynthates. Formation of the secondary cell wall (SCW) and lignification occurs in all xylem cell types. Vessel elements and fibers both undergo programmed cell death (PCD), with death in vessel elements occurring earlier than in fibers (Courtois-Moreau et al., 2009), while ray cells remain alive for several years (Nakaba et al., 2012). Distinct markers defining sub-stages of the differentiation processes are currently not available.

Transcriptomics and genetic analyses have been used to enhance our understanding of the underlying molecular mechanisms of secondary growth using *Arabidopsis thaliana* as a model system (Ruzicka et al., 2015; Fukuda, 2016). However, the small size and limited extent of secondary growth of *A. thaliana* severely limit the spatial resolution at which SCW formation and vascular development can be assayed. Additionally, the xylem fibers of *A. thaliana* do not fully mature but stay alive even during prolonged growth, at least in greenhouse conditions (Bollhöner et al., 2012). Use of woody tree systems, such as *Populus spp.,* overcomes these limitations and transcriptomic analyses in such species have enhanced our understanding of the molecular mechanisms underlying cambial growth and wood formation. The first study of the wood formation transcriptome was performed by Hertzberg et al. (2001) using cDNA microarrays covering less than 10% of the *Populus* gene space (2,995 probes). This study utilized tangential cryosectioning, as first described in Uggla et al. (1996), to obtain transcriptomes from different stages of wood development. Subsequently, a number of similar studies were performed using microarrays with higher gene coverage (Schrader et al., 2004; Aspeborg et al., 2005; Moreau et al., 2005; Courtois-Moreau et al., 2009) and, recently, RNA-Sequencing (RNA-Seq) (Immanen et al., 2016). However, earlier transcriptome studies of cambial growth and wood formation in *Populus* were limited to specific wood forming tissues using supervised approaches, i.e. samples were collected on the basis of visual anatomical assessment during sectioning.

The availability of RNA-Seq methodologies enables transcriptome profiling with higher dynamic range than was previously possible. Moreover, RNA-Seq does not require prior knowledge of gene models and can be used to identify effectively all transcribed loci in a sample (e.g. Goodwin et al. (2016)). The base-pair resolution of RNA-Seq additionally enables differentiation of expression from transcripts arising from highly sequence-similar paralogous genes or gene-family members. This is particularly pertinent for studies within the *Salicaceae* family, which underwent a relatively recent whole genome duplication (WGD) event, with about half of all genes still being part of a paralogous gene pair in the *P. trichocarpa* reference genome (Sterck et al., 2005; Tuskan et al., 2006; Goodstein et al., 2012). Concomitantly, tools for analyzing expression data, such as co-expression networks, represent increasingly powerful approaches for identifying central genes with important functional roles (e.g. Mutwil et al. (2011); Netotea et al. (2014)).

Here, we employed RNA-Seq to assay gene expression across wood-forming tissues in aspen (*Populus tremula*). Cryosectioning was used to obtain a continuous sequence of samples extending from differentiated phloem to mature xylem with high spatial resolution (25-28 samples per replicate). This enabled a continuous and unsupervised analysis of transcriptional modules, which were assigned developmental context using functionally characterized genes. We generated expression profiles for 28,294 previously annotated genes in addition to *de novo* identification of 78 protein-coding genes and 567 long intergenic noncoding RNAs. Co-expression network analysis was used to identify the 41 most central transcriptional modules representing major events in cambial growth and wood formation. Several modules could be related to discrete and well described processes such as cell division, cell expansion and SCW formation, while many other modules comprised genes with no characterised function. The high spatial resolution of the data also enabled discovery of potentially novel roles of well-characterised genes involved in cellulose and xylan biosynthesis, regulators of xylem vessel and fiber differentiation and lignification. A majority of paralogs derived from the Salicaceae WGD displayed distinctly different expression profiles across wood formation. However, we found evidence indicating that paralogs with conserved expression may have been retained to achieve high expression during SCW formation. The data and co-expression network are publically available in the AspWood (Aspen Wood, http://aspwood.popgenie.org) interactive web resource, which enables genomics analysis of cambial growth and wood formation.

## Results and Discussion

### The AspWood resource: high-spatial-resolution expression profiles for secondary growth

We have developed the AspWood resource, which contains high-spatial-resolution gene expression profiles across developing phloem and wood forming tissues from four wild-growing aspen (*P*. *tremula*) trees. The data was produced using RNA Sequencing (RNA-Seq) of longitudinal tangential cryosections (Uggla et al., 1996; Uggla and Sundberg, 2002) collected from a small block dissected from the trunk of 15-m high, 45-year-old clonal aspen trees during the middle of the growing season (Table S1). We pooled sections (each 15?m thick) into 25 to 28 samples for each replicate tree. The pooling was based on the estimated tissue composition from anatomical inspection during sectioning (Figure 1A and Table S2), with the aim of sequencing single sections across the cambial meristem, pools of three sections across expanding and SCW forming xylem, and pools of nine sections across maturing xylem. The last sample pool included the annual ring from the previous year. Mature phloem sections were pooled into a single sample. We mapped the RNA-Seq reads to the *P. trichocarpa* reference genome (Tuskan et al., 2006; Wullschleger et al., 2013), and classified 28,294 in the current annotation as expressed (Table S3). In addition, we identified novel protein coding genes, long intergenic non-coding RNAs (lincRNAs) and other gene fragments with undetermined coding potential, of which 78, 567 and 307, respectively, were expressed (Table S4, S5 and S6). Hence these data expand the *P. trichocarpa* annotated gene space with several new transcribed loci. The transcriptome showed high diversity across the entire sample series (Figure S1), with similar numbers of genes observed in all samples, both when considering all expressed, annotated genes (range 21,592 to 25,743 genes) or when considering those accounting for 90% of the expression (range 9,181 to 13,420 genes).

**Figure 1.**
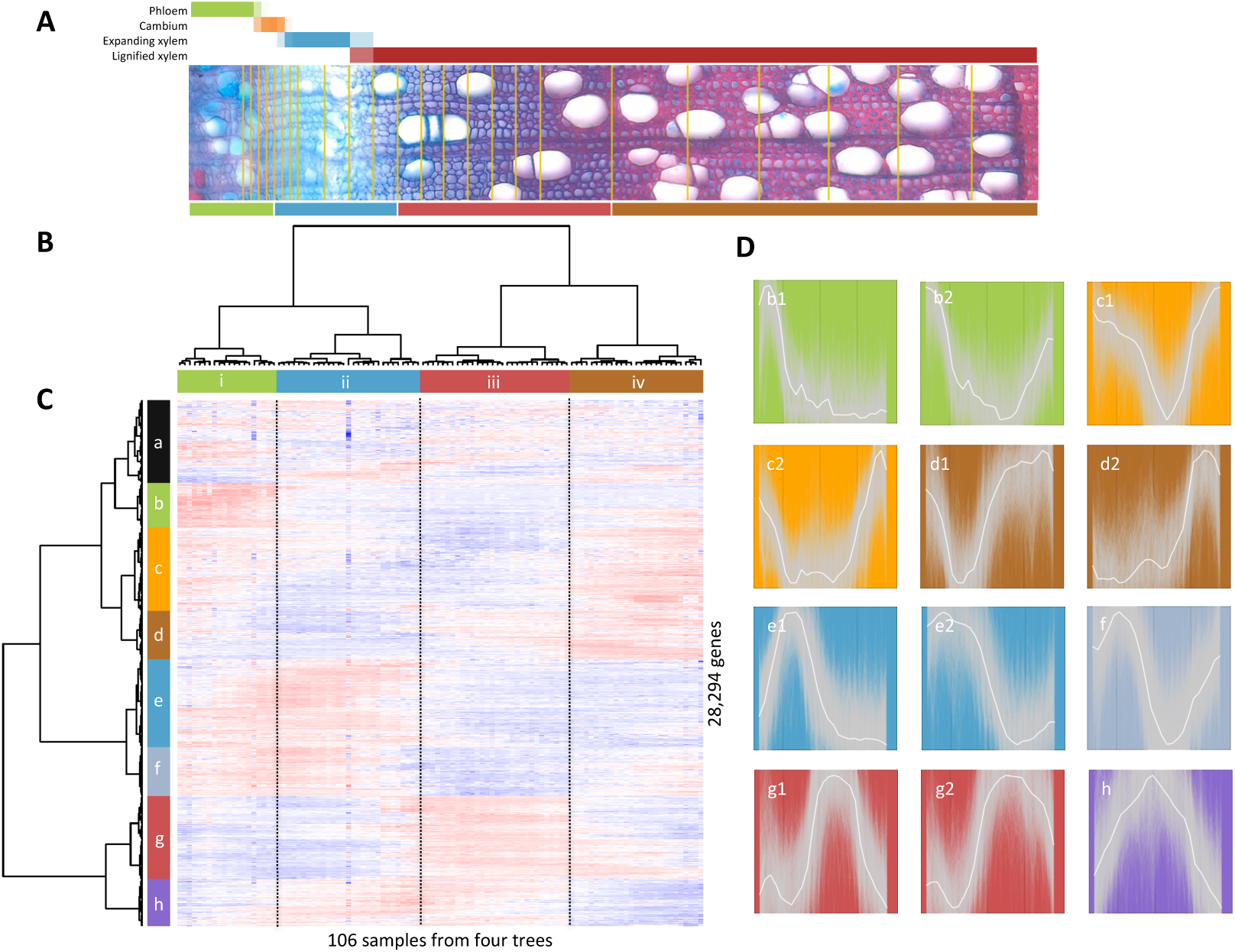
Hierarchical clustering of samples and genes across developing xylem and phloem tissues. (A) Transverse cross-section image from one of the sampled trees (tree T1). The strategy for pooling the samples for RNA-Seq is visualized by overlaying the samples on the section (positions on the section are approximated). The color bar below the image shows four sample clusters identified by hierarchical clustering (see B). The color bar above the image shows the estimated tissue composition for each sample. (B) Hierarchical clustering of all 106 samples from the four replicate trees using mRNA expression values for all expressed genes. The four main clusters are indicated with colors. (C) Heatmap describing hierarchical clustering of the 28,294 expressed annotated genes using mRNA expression values for all samples. Expression values are scaled per gene so that expression values above the gene average is shown in red, and below average in blue. Eight main clusters have been assigned colors and are denoted a to h. (D) Average expression profiles in tree T1 for each gene expression cluster and distinct sub-clusters (solid white lines). The expression profiles of all individual genes assigned to each cluster is shown as grey lines in the background.

We have made the AspWood data available as an interactive web resource (http://aspwood.popgenie.org). enabling visualization and exploration of the expression profiles and providing the scientific community with the opportunity to build and test hypotheses of gene function and networks involved in cambial growth and wood formation.

### Three transcriptome reprogramming events define cambial growth and wood formation

To provide an overview of the expression dataset, we performed an unsupervised hierarchical clustering analysis of all samples and annotated genes. This identified four major sample clusters (denoted ***i***, ***ii***, ***iii*** and ***iv***) defined by three distinct transcriptome reprogramming events (Figure 1AB, Figure S2, Table S2). These three events were assigned developmental context using expression patterns of well-characterized genes that we designate as markers for phloem differentiation (homolog of SUS6), cambial activity (*PtCDC2*), radial cell expansion (*PtEXPA1*), SCW formation (*PtCesA8-B*) and cell death (homolog of *BFN1,* see Figure S3 for profiles and documentation), which were further supported by anatomical data (Figure 1A). The first reprogramming event (***i***/***ii***) occurred between phloem and xylem differentiation, in the middle of the dividing cambial cells. The second event (***ii***/***iii***) marked the end of cell expansion and the onset of SCW formation. The final event (***iii***/***iv***) marked the end of SCW deposition, demonstrating that the late maturation of xylem cells should be considered a defined stage of wood development with a characteristic transcript-ome. We have indicated these three transcriptome reprogramming events as reference points (vertical dashed lines] in all expression profiles shown here and in the AspWood resource (e.g. Figure 2).

**Figure 2.**
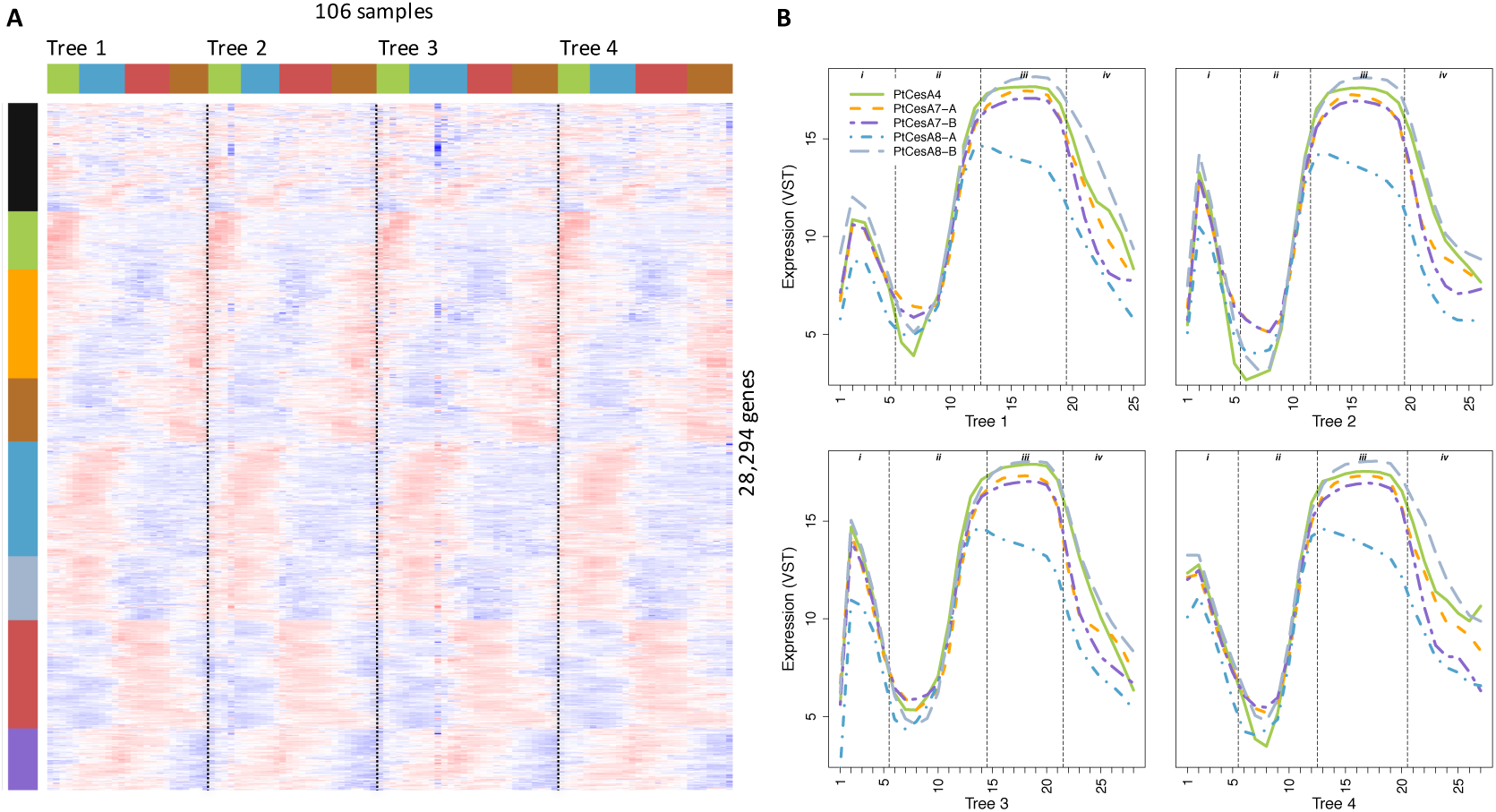
(A) Hierarchical clustering of genes across developing xylem and phloem tissues with samples ordered according to sampling order for each of the four replicate trees. The color bars indicate sample and gene clusters (see Figure 1). (B) Expression profiles of the secondary cell wall CesA genes in all four trees (T1-T4). PtCesA4 (Potri.002G257900), PtCesA 7-A (Potri.006G181900), PtCesA7-B (Potri.018G103900), PtCesA8-A (Potri.011G069600) and PtCesA8-B (Potri.004G059600).

We further used the hierarchical clustering to group genes, dividing the transcriptome into eight major gene expression clusters denoted ***a*** to ***h*** (Figure 1C, Table S7]. The expression clusters were highly reproducible in the four replicate trees (Figure 2], demonstrating that there is tight genetic control of the transcriptome throughout cambial growth and wood formation, even in a natural forest setting. With the exception of cluster ***a***, the gene expression clusters, or sub-clusters, had distinct average expression profiles (Figure 1D). These profiles revealed the major gene expression patterns underlying the three major transcriptome reprogramming events identified from the sample clustering.

### Co-expression network analysis reveals a continuum of transcriptional modules across cambial growth and wood formation

To identify potential novel regulators of secondary growth, we constructed a coexpression network (see Methods). The network connected pairs of genes with high normalized co-expression (Z-score > 5) and included 14,199 of the 29,246 expressed genes. Genes were then ranked according to their centrality in the coexpression network, i.e. the number of co-expressed genes in their network neighborhoods (Table S8). Transcription factors (TFs) were more central in the network than other genes (P = 2e-5), and also had more sample-specific expression profiles (i.e. pulse-like, narrow profiles with expression restricted to a few consecutive samples, see Methods, P < 2e-16). Notably, several of the *de novo* identified protein coding and lincRNA genes were highly central in the coexpression network (Table S8) indicating that they may serve important functional roles in cambial growth and wood formation in aspen (Table S8). However, compared to previously annotated genes, a higher proportion of novel genes were not integrated into the network (~30% vs. ~50%), which might reflect their evolutionary recent origin (i.e. low conservation, Table S8), as suggested by Zhang et al. (2015). Such genes represent potential species- or clade-specific adaptations and regulatory mechanisms.

To categorize the transcriptionally regulated biological processes in cambial growth and wood formation, we utilized the co-expression network to identify representative network modules (NMs) containing non-overlapping sets of genes that were highly co-expressed with the most central genes in the network (Figure 3). This unsupervised analysis relied only on the expression data and not on anatomical annotations or genes with known roles and thus represents an unbiased description of the central biological processes underlying cambial growth and wood formation. We assigned putative biological functions to the 41 NMs containing at least 20 co-expressed genes by performing Gene Ontology (GO) and Pfam (Protein family) enrichment analyses (Table S9, Figure 3). The NMs provided a detailed view of the transcriptional program underlying cambial growth and wood formation, revealing a far more fine-grained modularity than was apparent from previous transcriptional studies. Noticeably, the different NMs had distinct spatial expression domains that were distributed along the entire sample series from phloem to the annual ring border. This indicates that anatomical and histochemical markers that are often used to characterize phloem and xylem development are a manifestation of a continuum of transcriptional modules. We next used these NMs to provide developmental context to the three major transcriptome reprogramming events identified by the hierarchical clustering analysis (Figure 1).

**Figure 3.**
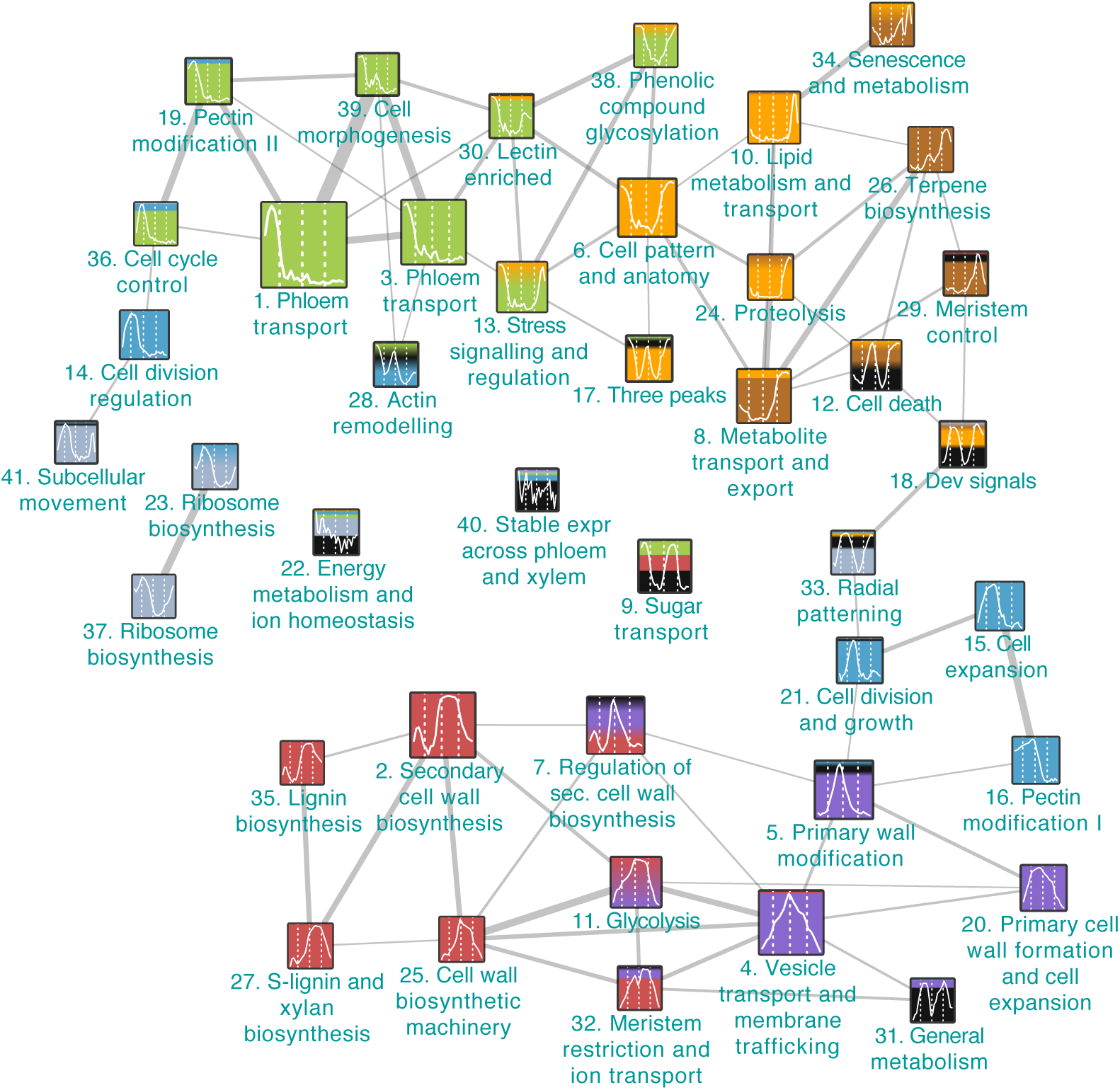
A modular version of the co-expression network. Genes with representative expression profiles were identified in the co-expression network (at a Z-score threshold of 5) by iteratively selecting the gene with the highest centrality and a co-expression neighborhood not overlapping with any previously selected genes’ neighborhood. Only annotated genes and positively correlated co-expression links were considered (i e Pearson correlation > 0). The selected genes and their co-expression neighborhoods (network modules) were represented as nodes in a module network, and given descriptive names (Table S9). The nodes are colored according to the hierarchical clusters in Figure 1, and reflect the proportion of genes in each network module belonging to the different hierarchical clusters The nodes were linked if the neighborhoods overlapped at a Z-score threshold of 4. Link strengths are proportional to the number of common genes. Overlaps of fewer than five genes were not represented by links, and only the 41 network modules with at least 20 genes were displayed.

We observed that genes within several NMs had expression profiles associated with a single developmental process (17 NMs with single peaks). NM1 (*Phloem transport,* Figure 3) contained genes associated with phloem identity and differentiation, including homologs of *ALTERED PHLOEM DEVELOPMENT (APL)* (Bonke et al., 2003), *CLAVATA3/ESR-RELATED 41* (*CLE41*) (Hirakawa et al., 2008; Etchells and Turner, 2010) and *KANADI (KAN)* (Emery et al., 2003; Ilegems et al., 2010), and was enriched for genes involved in *sucrose metabolism* (see Table S9 for enrichment details and Table S10 of unique gene identifiers), including homologs of *SUCROSE SYNTHASE* (*SUS*) *5* and *6,* which are known to be phloem expressed in *A. thaliana* (Barratt et al., 2009). Genes in this module had a distinct expression peak in the beginning of the sample series, with expression almost completely confined to sample cluster *i* (Figure 1), supporting functional roles in phloem differentiation and activity. NM14 *(Cell division regulation,* Figure 3) was enriched for the Pfam domain *cyclin,* known to be associated with the progression of the cell cycle, as well as other cell division regulatory genes like *CELL DIVISION CONTROL 2* (*PtCDC2*) (Espinosa-Ruiz et al., 2004). Expression of genes in this module peaked where the hierarchical clustering indicated the first transcriptome reprogramming event (***i****/****ii***), thus marking the cambium. NM15 (*Cell expansion,* Figure 3) was enriched for the Pfam domain *pectate lyase,* and included the alpha-expansin *PtEXPA1,* which has a demonstrated role in xylem cell expansion (Gray-Mitsumune et al., 2004; Gray-Mitsumune et al., 2008), and the xylem-specific pectate lyase *PtPL1-27* (Biswal et al., 2014). The expression of this module was specific to sample cluster ***ii***, consistent with genes in this module functioning in xylem cell expansion. Finally, NM4 (*Vesicle transport and membrane trafficking,* Figure 3) was enriched for genes involved in *vesicle-mediated transport,* and included genes encoding components of the secretory pathway and vesicle transport (e.g. SNARE-like, clathrin and coatomer). This module had an expression profile spanning the entire sample series, but with a clear expression peak where the hierarchical clustering indicated the second transcriptome reprogramming event (***ii****/****iii***). This module points to the importance of the secretion machinery at this stage of wood formation. Taken together, these modules can be used to identify novel candidate regulators of cambial activity and cambial derivative expansion and differentiation, in addition to providing novel marker genes associated with the fine-scale expression programs active during secondary growth.

We found that other processes of vascular differentiation were associated with several different expression peaks (24 NMs with more than one peak). Examination of both the hierarchical clusters and NMs identified co-expressed gene sets with biphasic expression patterns (i.e. profiles with two peaks), likely indicating that the same biological processes represented by these regulatory modules were expressed in two population of cells during, for example, both phloem and xylem formation. The phloem-xylem biphasic expression pattern was exemplified by NM2 (*Secondary cell wall biosynthesis*), NM27 (*S-lignin and xylan biosynthesis*) and NM35 (*Lignin biosynthesis*) (Figure 3), which included SCW *CesAs,* xylan and lignin biosynthesis genes. These modules had expression profiles with a low peak in sample cluster ***i*** and a high, broad peak in sample cluster ***iii***, marking the biosynthesis of SCWs in both phloem fibers and xylem cells. NM7 (*Regulation of SCW biosynthesis,* Figure 3) contained the three TFs *PtMYB20* (McCarthy et al., 2010), *PtMYB3* and *PtMYB21* (Zhong et al., 2013), homologous to A *thaliana MYB46* and *MYB83,* which can induce ectopic SCW formation. The module was characterized by a narrow expression peak marking the initiation of SCW formation in the xylem and a lower narrow expression peak on the phloem side, marking phloem fiber formation prior to visual differentiation of these phloem cells into fibers. This expression pattern is consistent with this module including regulatory switches that induce the broader expression profiles of the structural genes in modules such as NM2, 25, 27 and 35, and is an example of the finding that TFs had significantly narrower expression domains than other genes. Finally, NM34 *(Senescence and metabolism,* Figure 3) was enriched for the Pfam domain *late embryogenesis abundant protein,* and included a homolog of *BIFUNCTIONAL NUCLEASE I* (*BFN1*), which has been implicated in the post-mortem destruction of dead xylem vessel nuclei (Ito and Fukuda, 2002). This module had two narrow expression peaks, one late in sample cluster ***iii***, and the second late in sample cluster *iv.* The decrease in expression of these two peaks coincides with the loss of viability in vessels elements and fibers, respectively (Table S2), thus marking the border of living vessel elements and fibers.

Taken together, the NMs provided an unbiased framework enabling localisation of biological processes during cambial growth and wood formation, and revealed a greater complexity than had been realized on the basis of previous, lower spatial resolution. While many of the NMs could be associated with known processes, several remained poorly characterized and warrant future attention. A notable feature of many of the network modules, and in particular the uncharacterized ones, was the presence of two or three peaks of expression, possibly representing biological processes active in different cell types or at different stages during the differentiation of a specific cell type. The most central gene(s) from each NM represent a novel resource of markers for the transcriptional activity of the associated NM, and in cases where a NM can be assigned a biological function, as markers for those biological processes.

### Similarities in the regulation of early vascular differentiation in primary and secondary meristems

To identify similarities between the regulation of primary and secondary meristems during vascular development we examined the expression profiles of genes homologous to known regulators of early vascular differentiation in *A. thaliana* primary meristems. The two plasma membrane-associated proteins OCTOPUS (OPS) and BREVIS RADIX (BRX) are expressed in procambium and protophloem precursors, respectively, and are required for the correct specification of protophloem cells (Scacchi et al., 2010; Truernit et al., 2012). In sieve tube element precursors, the CLAVATA3/EMBRYO SURROUNDING REGION 45 (CLE45) peptide inhibits early protophloem development by signaling through the LRR-RLK receptor BARELY ANY MERISTEM3 (BAM3) (Depuydt et al., 2013; Rodriguez-Villalon et al., 2014). In support of these genes having a similar role during secondary vascular development, we identified that the closest homologs of *OPS (PtOPSs,* Table S10) were highly expressed across the cambium, whereas genes homologous to *BRX, CLE45* and *BAM3* were highly expressed in the differentiating phloem (sample cluster ***i***, Figure 4A). Another key gene in regulating phloem development encodes the MYB transcription factor ALTERED PHLOEM DEVELOPMENT (APL) (Bonke et al., 2003). APL acts as a positive regulator of phloem identity in *A. thaliana,* and *apl* plants show ectopic xylem formation where phloem cells normally form (Bonke et al., 2003). *APL* and the *APL* targets *NAC45/86* have been shown to be highly expressed in differentiating primary phloem (Furuta et al., 2014), and here we show that they are also expressed in differentiating secondary phloem (Figure 4B). This suggests that components regulating primary phloem development in *A. thaliana* roots function similarly during secondary phloem development in tree stems.

**Figure 4.**
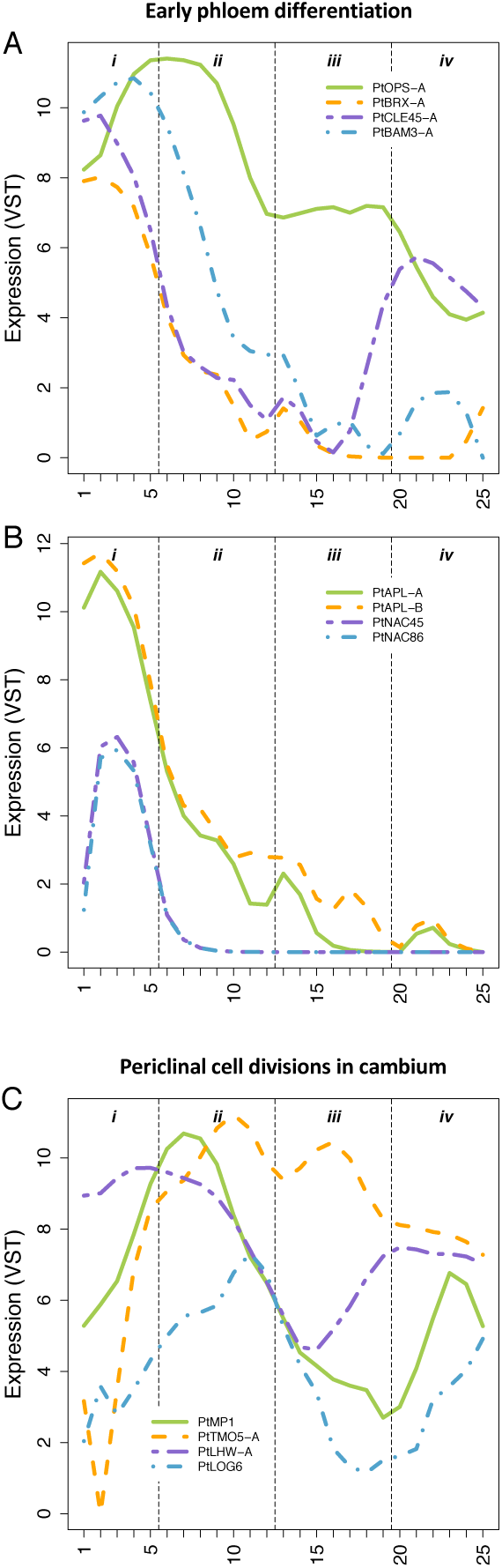
Similarities in expression patterns for regulators of early vascular differentiation between primary and secondary growth. (A) Expression profiles for the homologs of OPS, BRX, CLE45 and BAM3. PtOPS-A was highly expressed in the cambium while PtBRX-A, PtCLE45-A and PtBAM3-A were highly expressed in the secondary phloem (B) Expression profiles for the homologs of APL and NAC45/86. PtAPL and PtNAC genes were highly expressed in the secondary phloem. (C) Regulators of periclinal cell divisions. Expression profiles for genes homologous to MP, TMO5, LHW and LOG4 genes. High expression levels for all genes overlap in the expanding xylem. For the complete list of genes, see Table S10.

Establishment of vascular tissues in the embryo and the periclinal cell division activity of the root procambium in *A. thaliana* is regulated through the basic helix-loop-helix (bHLH) transcription factors TARGET OF MONOPTEROS 5 (TMO5) and LONESOME HIGHWAY (LHW) acting downstream of auxin signalling (Schlereth et al., 2010; De Rybel et al., 2013). TMO5 and LHW function as heterodimers, and co-accumulation of their transcripts in xylem precursor cells in root meristem is necessary and sufficient to trigger periclinal cell divisions of the adjacent procambium cells (De Rybel et al., 2013). This non-cell autonomous stimulation of cell division activity appears to act through local biosynthesis of cytokinin by the LONELY GUY3 (LOG3) and LOG4 (De Rybel et al., 2014; Ohashi-Ito et al., 2014), which acts as a mobile signal to promote periclinal cell divisions in the procambium. We found that although the closest homologs of the *TMO5* and *LHW* genes (Table S10) had complex expression patterns in the wood-forming zone of aspen, high expression of both genes intersected in the xylem expansion zone (sample cluster ***ii***, Figure 4C). Likewise, the *Populus LOG3/4* homolog *PtLOG6* (Immanen et al., 2013) was highly expressed in this region. Genes similar to *MONOPTEROS* (*MP/ARF5*), encoding an upstream regulator of the TMO5-LHW dimer (Schlereth et al., 2010), were expressed both in the cambium and in the expanding xylem. Taken together the expression profiles of the *Populus* homologs suggests that a bHLH complex involving similar components to that active in *A thaliana* roots likely regulate periclinal cell divisions during cambial growth in trees. Moreover, as in *A. thaliana* roots, the expression of *LOG* genes suggests a movement of cytokinins to stimulate cell divisions, in this case from the expanding xylem to cambial cells.

### Primary ceM-waM-polysaccharide biosynthetic genes continue to be expressed during secondary cell wall deposition in xylem tissues

In *Populus* xylem, primary and secondary cell wall layers have very different polysaccharide composition (Mellerowicz and Gorshkova, 2012). The PCW layers are abundant in pectins, such as homogalacturonan (HG) and rhamnogalacturonan I (RG-I), and they contain hemicelluloses such as xyloglucan, arabinoglucuronoxylan and mannan. In contrast, the SCW layers are rich in glucuronoxylan, and contain only small amounts of mannan. Cellulose is an important component in both layers, but the proportion of cellulose is significantly higher in SCW layers. These differences indicate that the cell wall polysaccharide biosynthetic machinery undergoes significant rearrangement during primary-to-secondary wall transition. The spatial resolution of AspWood enabled us to characterize changes in the transcriptome that occurred concurrently with this shift.

We identified the *Populus* genes encoding glycosyl transferases involved in the biosynthesis of HG (Atmodjo et al., 2011), RG-I (Harholt et al., 2006; Liwanag et al., 2012), RG-II (Egelund et al., 2006), and xyloglucan (Cavalier and Keegstra, 2006; Zabotina et al., 2008; Chou et al., 2012) (Table S10). As expected, the majority of these genes were highly expressed in primary walled cambial and radially expanding tissues (sample cluster ***ii***, Figure 5A). Interestingly, although expression peaked before the onset of SCW deposition, a majority were also significantly expressed during SCW deposition (sample cluster ***iii***), suggesting that there may be continuous biosynthesis of pectin and xyloglucan during SCW biosynthesis. The incorporation of these polymers could still occur in the primary wall layer, as suggested for xyloglucan in developing aspen fibers based on immunolabelling localization patterns (Bourquin et al., 2002). Moreover, transcripts of many enzymes involved in pectin metabolism such as pectate lyases and pectin methyl esterases (PMEs) were abundant in the SCW formation zone. Also transcripts of xyloglucan endotransglycosylases (XETs) were abundant in this zone, in agreement with previous reports of XET activity (Bourquin et al., 2002). This indicates a continuous metabolism of pectin and xyloglucan during SCW formation. Interestingly, the pectin biosynthetic genes *PtGAUT1-A, PtARAD1-A, PtGALS2-A* and *PtGALS2-B* had a distinct expression peak in mature xylem (sample cluster ***iv***, Figure 5A). The expression peaks of these genes may correspond to the biosynthesis of protective and isotropic layers deposited as tertiary wall layers after the deposition of SCW in the contact and isolation ray cells, respectively (Fujii et al., 1981; Murakami et al., 1999).

**Figure 5.**
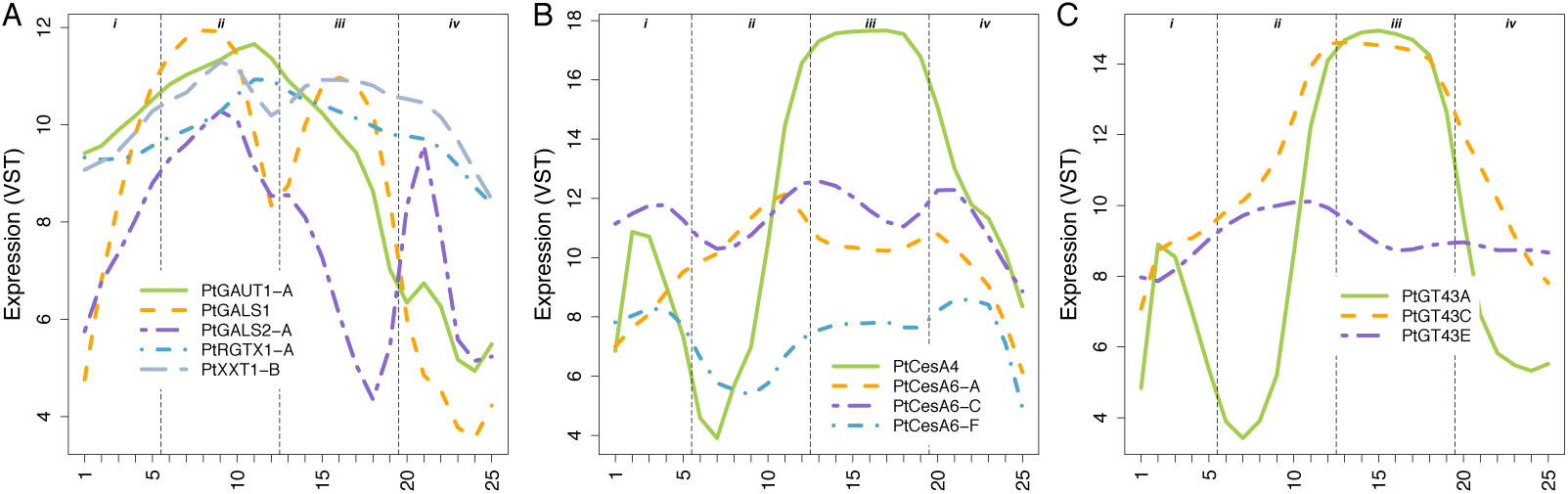
(A) Expression profiles for pectin and xyloglucan biosynthetic genes. All genes are highly expressed during primary wall biosynthesis, but also at later stages during xylem development i) representative expression for homogalacturonan biosynthesis genes (illustrated by PtGAUT1-A), ii) expression pattern of PtGALS1, iii) representative pattern of other RG-I biosynthesis genes (represented by PtGALS2-A), and iv) representative expression pattern for three of the putative xyloglucan biosynthesis genes (illustrated by PtXXT1-B). For a list of identified putative pectin and xyloglucan biosynthesis genes see Table S10. (B) Expression patterns for CesA genes. i) Members responsible for cellulose biosynthesis in the secondary wall layers are all induced in SCW biosynthesis zones in the xylem, and phloem (illustrated by PtCesA4) ii) Members classified as primary wall CesAs typically peak in primary wall biosynthesis zone, but are also highly expressed during later stages of xylem differentiation, (illustrated by PtCesA6-A), iii-iv) and some members even peak during these later stages (illustrated by PtCesA3-C, and 6-F). For a complete list of putative CesA genes see Table S10. (C) The GT43 gene family responsible for xylan biosynthesis comprise three clades, A/B, C/D and E, each having different expression here illustrated by i) PtGT43A, ii) PtGT43C and iii) PtGT43E. Expression profiles support the hypothesis that PtGT43A/B and PtGT43C/D are members of secondary wall xylan synthase complex, whereas PtGT43E and PtGT43C/D are members of the primary wall xylan synthase complex. For a complete list of genes co-regulated with the three clades of GT43 genes, see Table S10.

*A. thaliana* SCW cellulose synthase complexes contain equimolar trimers of CesA4, 7 and 8 (Hill et al., 2014), and *Populus* homologs forming similar complexes have been identified (Song et al., 2010). In AspWood, the corresponding transcripts showed a very specific expression pattern, marking the onset and end of SCW biosynthesis in the xylem (sample cluster ***iii***), as well as fiber differentiation in phloem (sample cluster ***i***, Figures 2B and 5B). The other *CesA* genes, homologous to *A. thaliana* “primary wall *CesAs”* (Kumar et al., 2009), showed varied expression patterns, with the majority highly expressed in the cambial and radially expanding tissues (Figure 5B), consistent with their role in cellulose biosynthesis in PCWs. Interestingly, there were also PCW *CesAs* that showed similar expression patterns as pectin and xyloglucan biosynthesis genes, with high transcript abundances during SCW biosynthesis, bringing into question their PCW specificity. A functional role for PCW CesAs during SCW biosynthesis is supported by the presence of protein complexes containing both PCW and SCW CesAs in xylem cells synthesizing SCW layers (Song et al., 2010; Carroll et al., 2012). Thus, what are referred to as PCW *CesA* genes might have a more general role than previously thought, including the possibility that they enter into complexes with SCW CesAs.

Arabinoglucuronoxylan of PCW layers and glucuronoxylan of SCW layers have the same backbone of β-1,4-Xyl*p* residues, which is thought to be synthesized by a heteromeric xylan synthase complex in the Golgi (Oikawa et al., 2013). At least three interacting glycosyltranferases (GT) of this complex have been identified, i.e. one GT47 member (IRX10/10L) and two GT43 members (IRX9/9L and IRX14/14L) (Jensen et al., 2014; Urbanowicz et al., 2014; Mortimer et al., 2015; Zeng et al., 2016). In *Populus,* IRX9/9L function is performed by PtGT43A, B, or E, and IRX14/14L function by PtGT43C or D (Lee et al., 2011; Ratke et al., 2015). In AspWood, expression of *PtGT43A* and *B* was almost identical and closely resembled that of SCW CesAs, whereas expression of the genes encoding their interacting partners PtGT43C or D was slightly different, with higher expression in primary-walled tissues (Figure 5C). *PtGT43E* showed a very different expression profile, with a broad expression peak in primary walled cells. These observations support the recently suggested concept that separate primary and secondary xylan synthase complexes exist, similar to primary and secondary cellulose synthase complexes (Mortimer et al., 2015; Ratke et al., 2015), with PtGT43E (IRX9L) and PtGT43C/D (IRX14) forming the primary xylan synthase complex, and PtGT43A/B (IRX9) and PtGT43C/D (IRX14) forming the secondary xylan synthase complex. Distinction between the expression patterns of PtGT43A/B and PtGT43C/D in AspWood highlighted the high resolution of the data and demonstrated the potential for both discovering new genes involved in wood formation and refining the expression domain of known and novel genes.

Taken together, the expression profiles of the pectin, xyloglucan and cellulose PCW biosynthetic genes suggest that the primary wall biosynthetic program is not terminated at the onset of secondary wall biosynthesis. It rather continues in the background of the more spatially confined expression of SCW genes such as CesA 4/7/8 and PtGT43A/B. Indeed, when SCW deposition is terminated in long-lived cell types such as parenchyma cells or gelatinous-fibers in tension wood, a subsequently deposited layer containing pectin and xyloglucan can be observed in these cells that resembles the PCW layer (Fujii et al., 1981; Murakami et al., 1999; Mellerowicz and Gorshkova, 2012). Our analyses additionally revealed that some genes of the SCW biosynthesis program are also active during the period of PCW formation. This was exemplified by PtGT43C/D and its co-expression network neighborhood, which included many proteins involved in general cellular functions associated with cell wall biosynthesis, such as vesicle trafficking, transport, sugar nucleotide metabolism and general cell wall acetylation machinery (Table S10).

### Spatially separated expression of phenoloxidases may enable site and cell type specific lignification

Cell wall lignification results from the secretion of differently methoxylated 4-hydroxyphenylpropanoids, called monolignols, which are radically oxidized by cell wall resident phenoloxidases (laccases and peroxidases) to cross-couple the monolignols into an insoluble polyphenolic polymer in the cell wall (Boerjan et al., 2003; Barros et al., 2015). In *Populus,* 92 genes encoding monolignol bio-synthetic enzymes have been identified (Shi et al., 2010) (Table S10), of which 59 were expressed in AspWood (Figure 6A, Table S10). The lack of expression for many monolignol genes may suggests that these are involved in the biosynthesis of phenolic compounds other than wood lignin. About half of the expressed monolignol biosynthetic genes had a bi-phasic expression profile, with a low peak in the differentiating phloem (i.e. sample cluster ***i***) and a high peak in the maturing xylem (clusters ***iii***), and with relatively high expression during late xylem maturation (i.e. sample cluster ***iv***, Figure 6AB). However, some were expressed only in the phloem or in the late maturing xylem (Figure 6A).

**Figure 6.**
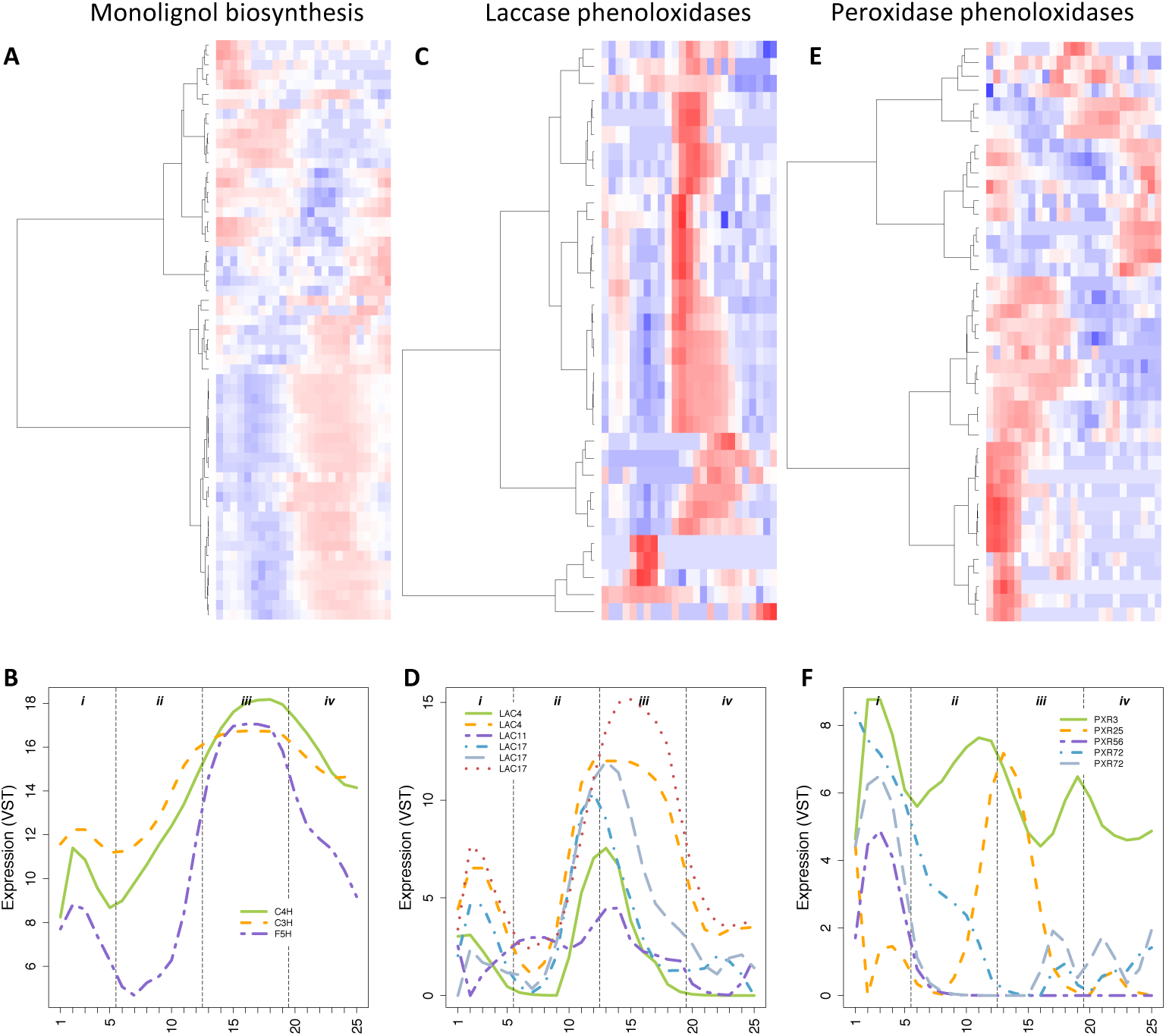
Expression profiles of genes involved in the lignification of xylem cells. (A) Hierarchal clustering and heatmap of the 59 monolignol biosynthesis genes expressed in AspWood. Expression values are scaled per gene so that expression values above the gene average are represented by red, and below average by blue. (B) Expression profiles of C4H (Potri.013G157900), C3H (Potri.006G033300) and F5H (Potri.005G117500) homologs. (C) Hierarchical clustering of the 34 expressed laccase phenoloxidases, and (D) expression profiles of representative genes: homologs of LAC4 (Potri. 001G248700, Potri. 016G112100), LAC11 (Potri. 004G156400) and LAC17 (Potri. 001G184300, Potri. 001G401300 and Potri. 006G087100). (E) Hierarchical clustering of the 42 expressed peroxidase phenoloxidases, and (F) expression profiles of representative genes: homologs of PXR3 (Potri.003G214800), PXR25 (Potri.006G069600), PXR56 (Potri.007G122200) and PXR72 (Potri.005G118700 and Potri.007G019300).

In *Populus,* 165 genes encoding phenoloxidases have been identified (Lu et al., 2013) (Table S10). Both peroxidases and laccases have been shown to be active in wood forming tissues of *Populus* and to be capable of polymerizing both mono-methoxylated (guaiacyl G) and bi-methoxylated (syringyl S) monolignols into lignin-like polymers *in vitro* (Christensen et al., 1998; Ranocha et al., 1999; Sasaki et al., 2004; Sasaki et al., 2008). More recently, laccases *LAC4/IRX12, LAC11* and *LAC17* (Zhao et al., 2013) were demonstrated to act redundantly during vessel element and fiber lignification. Lignin formation has also been shown to be dependent on the additive activities of multiple peroxidases, including *PXR2, PXR25, PXR71* and *PXR72* in *A. thaliana* xylem (Herrero et al., 2013; Shigeto et al., 2015) and the homolog of *PXR3* in *Populus* (Li et al., 2003). We found that 34 out 56 annotated laccases and 42 out of 109 peroxidases were expressed in AspWood (Table S10), and that these genes generally had narrower expression peaks than monolignol biosynthetic genes (p < 5e-5, Table S10, Figure 6CD). The highest expression of most laccases occurred at different phases of SCW formation (sample cluster ***iii***), with some also having a lower expression peak in the differentiating phloem (i.e. sample cluster ***i***). Only three peroxidases were co-expressed with laccases (Z-score threshold of five), with most peroxidases being expressed in the phloem and/or the cambium (i.e. sample clusters ***i*** and ***ii***) and in the late maturing xylem (i.e. sample cluster ***iv***, Figure 6EF). While experiments in *A. thaliana* have suggested that laccases and peroxidases act non-redundantly (Zhao et al., 2013), our data confirm that these enzymes have clearly separate expression domains in aspen, and suggest that laccases are primarily associated with lignification in the phloem and SCW formation zone while the peroxidases primarily are associated with lignification in the phloem, cambium and mature xylem.

### Transcriptional regulators of wood development

The differentiation of xylem vessel elements and fibers is regulated by NAC domain transcription factors (for reviews, see Zhong et al. (2006), Ruzicka et al. (2015) and Ye and Zhong (2015)). In *A. thaliana,* VASCULAR-RELATED NAC-DOMAIN 6 (VND6) and VND7 specify vessel element cell fate (Kubo et al., 2005; Yamaguchi et al., 2008), while VND1 to 5 act upstream of VND7 in formation of vessel elements (Endo et al., 2009; Zhou et al., 2014). Xylem fiber differentiation is mediated by SECONDARY WALL-ASSOCIATED NAC DOMAIN 1 (SND1) and NAC SECONDARY WALL THICKENING PROMOTING 1 (NST1) (Zhong et al., 2006), with the NAC domain transcription factors SND2 and 3 also contributing to fiber differentiation (Wang et al., 2013; Shi et al., 2017).

The *P. trichocarpa* homologs of *SND/VND* separate into five distinct phylogenetic clades (Figure 7A). Within a given clade, paralogs were typically highly co-expressed with one notable exception; members of the VND3-clade displayed similar expression patterns to either the *VND6-* or *SND2*-clade genes (Figure 7A-E). For members of the *VND6*-clade, expression peaked within the cell expansion zone (sample cluster ***ii***) and again further inwards, at the end of the SCW formation zone (sample cluster ***iii***, Figure 7E). Early expression of the *VND6-* clade genes is in line with a role as vessel element identity genes, with cell fate being specified at the onset of radial expansion. The continuous expression of VND6-clade genes during SCW deposition is consistent with the suggested role of these genes in SCW formation and cell death in vessel elements (Derbyshire et al., 2015). While one of the *Populus* VND7-clade paralogs was not expressed in our data, expression of the second VND7-clade paralog increased sharply in the SCW formation zone (sample cluster ***iii***, Figure 7D), overlapping with the inner peak of the *VND6* clade paralogs. The network neighborhood of the expressed VND7-clade paralog included a homolog of METACASPASE 9, which in *A. thaliana* has been shown to participate in the autolysis of xylem vessel elements (Bollhöner et al., 2013), as well as other genes (in total 77 genes at a Z-score threshold of three) suggesting that maximal expression of VND7, and the later peak in the expression profiles of the *VND6*-clade paralogs, marks the transition from the SCW to the maturation phase in xylem vessel elements. Thus, the spatial resolution of our sample series enabled us to define two clearly separated expression domains in vessel element differentiation.

**Figure 7.**
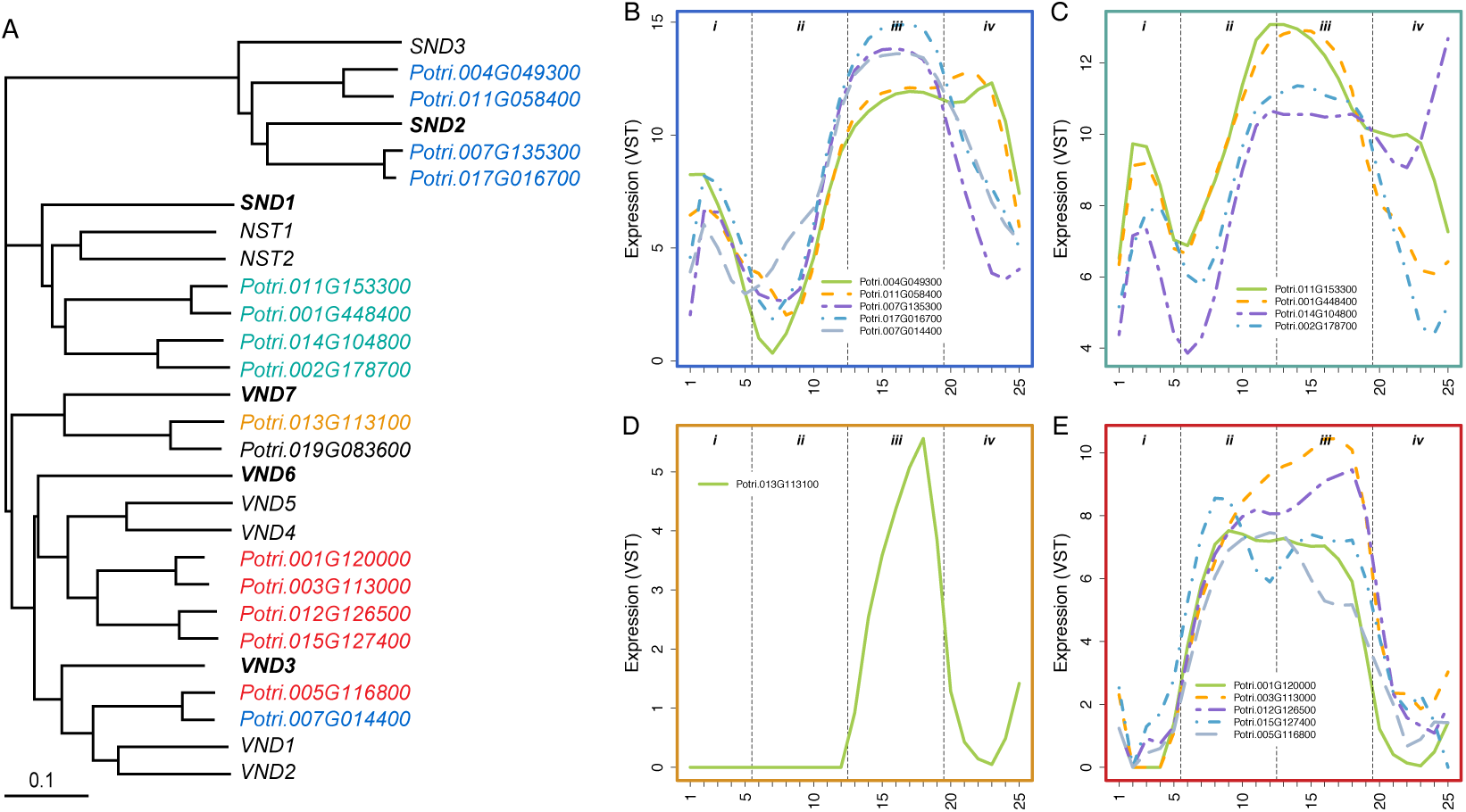
NAC domain transcription factors are expressed in distinct patterns corresponding to phylogenetic clustering. (A) Phylogenetic tree of Populus wood associated NAC-domain transcription factors with homologs in A. thaliana. Colors indicate co-expression. A. thaliana gene names in bold are used as clade names. (B) Expression profiles of Populus SND2 (Potri. 007G135300; Potri. 017G016700) and SND3 (Potri.004G049300; Potri.011G058400) homologs compared to the VND3 homolog with a similar expression profile (Potri.007G014400). (C) Expression profiles of Populus SND1 homologs (Potri.001G448400; Potri.002G178700; Potri.011G153300; Potri.014G104800). (D) Expression profile of the Populus VND7 homolog expressed in AspWood (Potri. 013G113100; Potri. 019G083600 is not expressed). (E) Expression profiles of Populus VND6 homologs (Potri. 001G120000; Potri. 003G113000; Potri. 012G126500; Potri.015G127400) compared to the VND3 homolog with a similar expression profile (Potri. 005G116800).

Compared to the VND6-clade genes (Figure 7E), expression of the *SND1*-clade genes was shifted slightly towards maturing xylem, but reached maximal values before the transition into the SCW formation zone (i.e. sample cluster ***iii***, Figure 7C). However, an almost perfect overlap with the SCW formation marker *PtCESA8-B* was found for the expression patterns of the *SND2* paralogs (Figure 7B). Unlike the *VND6*-clade paralogs, both *SND1* and *SND2* paralog expression increased gradually from the cambium towards the phloem in sample cluster *i*, indicating a role in SCW formation in phloem fibers. This suggests that the role of *SND1* and *SND2* paralogs is specific to fibers but that they do not discriminate between phloem and xylem. As was the case for VND6/7-clade genes, some of the SND1/2-clade genes also had two separate expression domains in the xylem, with the inner peak of SND paralogs coinciding with the loss of viability of fibers.

Taken together, NAC domain transcription factors involved in xylogenesis were expressed in four distinct domains. Early onset of expression of genes in both the *VND6* and *SND1* clades in the cell expansion zone is in support of their role in the specification of vessel elements and fibers, respectively. However, their expression remained high during the entire SCW phase (sample cluster ***iii***), consistent with roles connected to the deposition of wall polymers and/or *postmortem* autolysis.

We explored whether 110 direct targets of the four *SND1*-clade genes were coexpression neighbors in our network, including 76 genes identified as targets using transactivation assays and chromatin immunoprecipitation (ChIP) in protoplast-derived xylary tissue with a low degree of differentiation (Lin et al., 2013) and 34 genes (hereafter, non-ChIP targets) identified as transactivation targets based on different lines of experimental evidence (Ye and Zhong, 2015) (see Table S10 for all target genes). Seventy-six of these 110 putative direct targets of SND1 paralogs were expressed in AspWood, of which 68 (89%) formed a connected co-expression network with the four *SND1*-clade paralogs using a co-expression threshold of three (Figure S5). Interestingly, the majority of the ChIP targets were negatively correlated with the *SND1* paralogs (subcluster II and III in Figure S5), indicating that *SND1*-like transcription factors likely suppress transcription of these targets.

Having established that the co-expression network predicts previously characterised TF-target interactions, we subsequently analyzed network neighborhoods of high centrality TFs with no currently known functional role during wood formation (using Table S8). We focused on TFs in the hierarchical clusters ***g*** and ***h***, with expression profiles indicating a role in SCW formation (Figure 1). While the most central TF in cluster ***g*** has a known role in wood formation (homolog of *SND2*), the second-most central TF is homologous to Agamous-like 62 (*AGL62)*. This gene has an essential role in seed development in *A. thaliana,* where it is expressed exclusively in the endosperm (Kang et al., 2008). However, our data show that the homolog of this TF had a pronounced expression profile in wood formation. A GO enrichment test of the network neighborhood identified enrichment for *cellulose biosynthetic process* (p < 0.05, Z-score threshold of five). Interestingly, one of the *AGL62* paralogs in *Populus* has been linked to biomass yield in a GWAS study (Allwright et al., 2016). In cluster ***h***, the most central TF is also a known regulator PtMYB20 (homolog of MYB83), while the second-most central TF is uncharacterized (Potri.006G208100), with the network neighborhood being enriched for the GO category *cell wall organization or biogenesis.* These two examples demonstrate the power of using AspWood as a source for identifying novel regulators of cambial growth and wood formation for subsequent downstream biological characterisation.

### A majority of paralogs show differential expression during wood formation

In *Salicaceae,* a whole genome duplication (WGD) occurred relatively recently (58 million years ago, Dai et al. (2014)). WGDs are believed to be a major contributor to the evolution of novel function in genomes (reviewed in Hermansen et al. (2016)), and to contribute to species diversification. Gene duplicates resulting from WGDs, hereafter called paralogs, may evolve through different processes; a) sub-functionalization, where the genes retain different parts of the ancestral function, b) neo-functionalization, where one gene retains the ancestral function while the other evolves a new function, or c) nonfunctionalization, where one of the genes is eliminated by random mutations. These processes may act both on protein function and on gene expression, with regulatory divergence being particularly important for evolution of plant development (Rosin and Kramer, 2009). A previous microarray study in *Populus* covering 14 different tissues indicated that nearly half of the paralog pairs have diverged in their expression (Rodgers-Melnick et al., 2012). We used our RNA-seq data to map the regulatory fate of WGD derived paralogs in cambial growth and wood formation. Of the 9,728 paralogous gene pairs identified in the *P. trichocarpa* genome by sequence similarity and synteny (see Methods), 8,844 had at least one gene expressed in our data. Of these paralogs, 3,185 (36%) had diverged regulation, as defined by their presence in different expression clusters *and* by their expression correlation being below 0.5 (Figure 8A, Table S11A). An additional 1,930 paralogs (22%) had only one gene expressed in our data. These may represent cases of non-functionalization, or cases where one gene is expressed in another tissue or condition. Paralogs with diverged expression were often found to be enriched in spatially-adjacent cluster-pairs (e.g. present in clusters ***g*** and ***d***) (Figure 8A, Table S11B), but paralogs with more diverged expression were also observed. For example, 1,163 paralogs (17%) had negatively correlated expression profiles, of which 156 displayed mirrored profiles (correlation < −0.50, see e.g. Figure 8B). Our estimate of regulatory divergence can be considered conservative, representing cases where paralogs display distinctly different expression profiles across cambial growth and wood formation. However, the high spatial resolution of our dataset also enabled the identification of far more subtle differences, such as the previously reported differences in expression levels between *CesA8-A* and *CesA8-B* in secondary phloem and xylem (both located in cluster ***g***, Figure 8C (Takata and Taniguchi, 2015). While the processes of sub-, neo- or non-functionalization drive the divergence of paralog pairs, the gene dosage balance hypothesis may explain why some paralog pairs have retained similar expression profiles (Yoo et al., 2014). Interestingly, a high proportion of the paralog pairs with similar profiles were found among the 5% most abundant transcripts in our dataset (51%, p = 5e-92). In particular cluster ***g***, including genes expressed during SCW formation (sample cluster ***iii***), contained a substantially higher fraction of such highly expressed and regulatory conserved paralogs than other clusters (Figure 8D).

**Figure 8.**
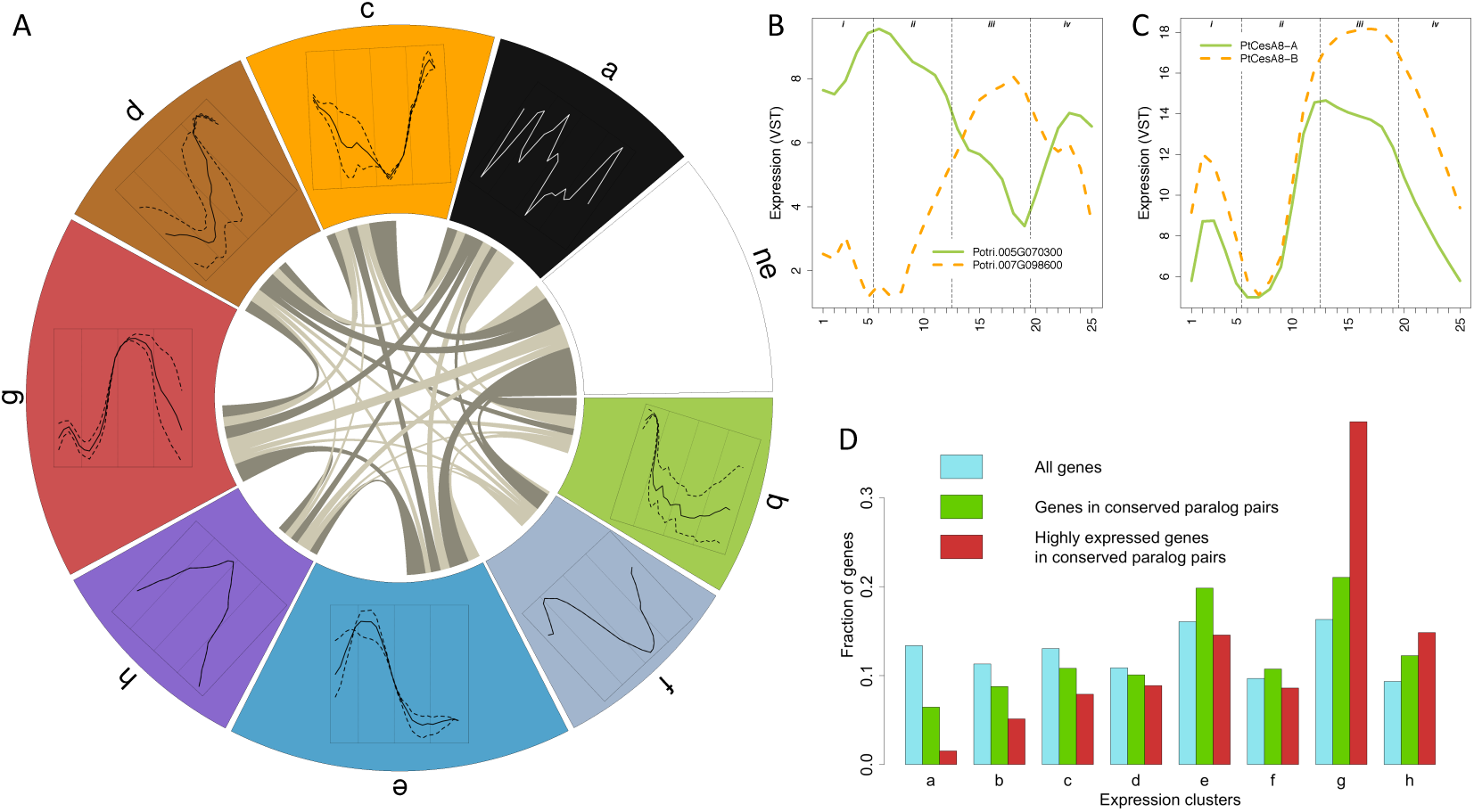
WGD paralog expression. (A) Circus plot of regulatory diverged and conserved paralogs. The clusters are ordered according to their peak of expression along the wood developmental gradient An additional cluster for genes not expressed (ne) in our data is also added. Each cluster occupies a share of the circle’s circumference proportional to the number of genes in that cluster belonging to a paralog pair. Paralogs expressed in two different clusters are shown by links. The width of a link is proportional to the number of paralogs shared between these clusters (i.e. diverged pairs). Only pairs in different clusters and with an expression Pearson correlation < 0.5 were considered diverged. Links representing more pairs than expected by chance (P < 0.0001) are colored in a darker tone. The share of a cluster without links represents the proportion of paralogs where both genes belong to that same cluster (i.e. conserved pairs). (B) Example of paralogs with highly diverged profiles (correlation = −0.78): FAAH (fatty acid amide hydrolase, Potri.005G070300 (cluster ***e***) and Potri.007G098600 (cluster ***g***)). (C) Previously published real-time PCR data of CesA genes showed that PtCesA8-B (Potri.004G059600) was expressed higher than PtCesA8-A (Potri.011G069600) in secondary phloem and xylem (Takata and Taniguchi, 2015). Although these genes have highly similar expression profiles (Pearson correlation = 0.93), and are therefore considered as conserved according to our definition (both belong to cluster ***g***), the data confirm the difference in expression levels and show that even more subtle regulatory divergence can be detected in the dataset. (D) The distribution of different gene sets among expression clusters: (blue) all 28,294 expressed, annotated genes in our data, (green) all 3,729 paralog pairs with conserved expression (i.e. in the same expression cluster), (red) 721 genes with high expression (among the top 5% most highly expressed genes in our data; 1,413 genes) and in paralog pairs with conserved expression.

This suggests that it has been advantageous for *Populus* to maintain higher levels of certain genes involved in SCW formation than could be achieved by a single ancestral gene copy. A second copy can also make the system more robust to perturbations.

### AspWood: a new reference resource for wood biology

The AspWood resource comprises high spatial resolution gene expression profiles of the protein-coding and non-coding transcriptome underlying cambial growth and wood formation in a model angiosperm forest tree, with sampling extending from the phloem across the cambium and wood forming tissues to the previous year’s annual ring. The associated web resource provides the scientific community with an interactive tool for exploring the expression profiles and their relatedness within the corresponding co-expression network. We used the network to identify a continuum of transcriptional modules covering the entire sample series, from which novel expression profiles that can serve as markers for biological processes active during cambial growth and wood development can be extracted. We also showed how the network can be used to identify validated targets of transcription factors acting both as activators and repressors, and demonstrated a strategy for identifying novel regulators of wood formation. Finally, we demonstrated that the resource enables the discovery of previously uncharacterised expression domains of well-studied genes, as well as the identification of expression differences in highly sequence-similar paralogs arising from the Salicaceae whole genome duplication. The AspWood resources represents a new reference resource for wood biology, and a natural starting point for designing functional experiments to further refine understanding of the transcriptional regulation of ligno-cellulose production in trees.

## Methods

### Sampling and RNA extraction

Wood blocks were collected from four independent naturally growing *P. tremula* clones (tree IDs T1, T2, T3 and T4) from Vindeln, north Sweden. Hand sections were taken from freshly collected stem material for analysis of cell viability using nitroblue tetrazolium (Courtois-Moreau et al., 2009). 15 micron thick longitudinal sections were cut using a cryo-microtome (Uggla et al., 1996) and stored at −80C. Cross-sections were taken during the process of sectioning and imaged with a light microscope. These images were used to characterize the different tissue types present. All cryosections were deemed to originate from one of four developmental zones: phloem, cambium, early/developing xylem and mature xylem. No obvious biotic or abiotic stresses were evident at the time of sampling and anatomically discernible developing tension wood was avoided. The total number of sections varied from 105 to 135 in the four different trees, and covered the entire current year’s growth.

Individual sections were thawed in QIAzol Lysis Reagent (Qiagen, Manchester, UK) and homogenised using a Retsch mixer mill. Total RNA and small RNA were extracted from section pools as indicated in Table S2. The miRNeasy Mini Kit (Qiagen, Manchester, UK) was used for extractions. For Total RNA, one elution of 35 μl was made. For small RNA, two elutions of 25 μl each were made and pooled.

Total RNA was quantified using a Nanodrop 1000 (Thermo Scientific, Wilmington, USA) and quality was assessed using the Agilent 2100 Bioanalyzer (Agilent Technologies, Santa Clara, USA) using Pico chips (Agilent Technologies, Santa Clara, USA). A 100 ng aliquot was used for amplification. If the volume of the 100 ng aliquot was larger than 5 μ1 it was dried using a speed-vac. In some cases, less than 100 ng total RNA was available, in which case the maximum possible amount was used. Total RNA was amplified using Ambion MessageAmp™ Premier RNA Amplification Kit (Ambion, Thermo Scientific, Wilmington, USA following manufacturer’s instructions. Using 100 ng of total RNA as starting material, we obtained 10-63 μg of amplified RNA (aRNA). aRNA was quantified using a Nanodrop and integrity checked using a Bioanalyzer with Nano chips (Agilent Technologies, Santa Clara, USA). 2.5 μg of aRNA for each sample was used for sequencing using 2×100 bp non-stranded reads on the Illumina HiSeq 2000 platform, as detailed in Sundell et al. (2015). Raw RNA-Seq data was uploaded to the European Nucleotide Archive (ENA, http://www.ebi.ac.uk/ena/): accession number ERP016242.

### RNA-Seq preprocessing

The RNA-Seq data was analyzed using a previously developed pipeline for quality control, read mapping and expression quantification (Delhomme et al., 2014). Briefly, we used FastQC for quality control (http://www.bioinformatics.babraham.ac.uk/projects/fastqc/), STAR for read mapping (Dobin et al., 2013), HTSeq for read counts (Anders et al., 2014) and DESeq for (variance stabilized) gene expression values (Love et al., 2014). The *P. trichocarpa* v3.0 genome and gene annotations were used (phytozome.org, v10) (Goodstein et al., 2012). Samples T4-05, T4-09, T4-20 and T4-28 showed low reads counts (Table S12A), while T3-17 was a clear outlier in a principal component analysis (PCA) of all samples (Table S12B). T4-28 was the last sample of T4 and was removed in subsequent analysis. The other four samples were replaced by the average expression values of the two flanking samples. PCA was performed using the R function *prcomp.*

To investigate whether the four trees were clonal replicates or not, a genotype test was performed using Single Nucleotide Polymorphisms (SNPs) in all genes from chromosome 1 in the *P. trichocarpa* genome assembly. RNA-Seq reads from the four replicates, as well as for four different genotypes from an independent experiment (control), were merged using samtools-1.3.1 mpileup (-d100000). SNPs were called using bcftools-1.3.1 call (-v -c). A PCA plot was created from the resulting variant call format file (vcf; Li et al. (2012)) using the R package SNPRelate-1.6.4 (Zheng et al., 2012). From the PCA plot it was clear that the four trees were clonal replicates (Figure S6).

We also developed a pipeline for annotating novel protein coding genes and long intergenic non-coding RNAs (lincRNA) using a combination of establish annotation tools. We used PASA (Haas et al. (2003), 2.0.3, default settings) to construct a reference database based on three assemblies: one *de novo* transcript assembly using the Trinity *de novo* pipeline (Grabherr et al. (2011), r20140717, settings: --min_kmer_cov 1 --normalize_reads --normalize_by_read_set) and two genome guided assemblies using Trinity (Grabherr et al. (2011), r20140717, settings: --genome --genome_guided_use_bam --genome_guided_max_intron 11000 --min_kmer_cov 1) and cufflinks (Trapnell et al. (2010), 2.2.1, settings: --l 24 library-type fr-unstranded-I 15000 --no-faux-reads). PASA reported 59 novel protein coding genes as well as 21,938 Expressed Sequence Tags (ESTs) assemblies with a CDS shorter than 40% of the mRNA length. To identify intergenic RNAs, we removed any region overlapping known *P. trichocarpa* genes (using bedtools (Quinlan and Hall, 2010), v2.19.1, settings: subtract-A). This retained 53 protein-coding genes and 13,995 ESTs. Potential frame-shift errors were identified using frameDP (Gouzy et al. (2009), 1.2.2, default settings) and 672 full length sequences, with CDSs longer than 40% of the mRNA length after correction, changed status to novel protein-coding genes. 816 ESTs with only a start coding, only a stop codon or neither a start nor a stop codon were classified as fragments. The remaining ESTs (12,507 assemblies) were classified as lincRNAs.

Genes were classified as expressed if the variance stabilized gene expression value was above 3.0 in at least two samples in at least three of the four replicate trees. Furthermore, we only considered novel genes longer than 200 bp (see e.g. Ulitsky (2016)). This resulted in 28,294 annotated genes, 78 novel proteincoding genes, 567 lincRNAs and 307 fragments with expression during aspen wood development.

### Expression analysis

Samples were clustered using Euclidean distance in the R (R-Core-Team, 2012) function *hclust.* Genes were scaled and clustered using Pearson correlation. Ward’s method was used in both cases. Dendrograms and heatmaps were generated using the R function *heatmap.2* in the *gplots* library. Samples were reordered within the dendrogram to best match the order in which they were sampled using the R function *reorder*. Functional enrichment was tested using Fisher’s exact test and false discovery rate corrected. Annotations were downloaded from Phytozome (Goodstein et al., 2012).

To group paralogous genes, the primary protein sequence of 41,335 *P. trichocarpa* genes (Phytozome,Goodstein et al. (2012)) were compared to each other by an all-against-all BLASTP following by a Markov Clustering algorithm (TribeMCL). Genomic homology was detected using i-ADHoRe 3.0 (Proost et al., 2012), using the following settings: alignment method gg4, gap size 30, cluster gap 35, tandem gap 10, q value 0.85, prob cutoff 0.001, anchor points 5, multiple hypothesis correction FDR and level 2 only true. Gene cluster pairs with significantly many shared paralog-pairs (i.e. with one gene in each cluster) were found by comparing the clustering to 10,000 randomly generating clusterings. A circus plot was created using the R function *circos* in the *circlize* library.

Co-expression networks were inferred using mutual information (MI) and context likelihood of relatedness (CLR,Faith et al. (2007)). The CLR algorithm computes a Z-score for each gene pair by using the MI value to all other genes as a null model. A co-expression network was constructed by linking all genes, including annotated genes, novel protein-coding genes and lncRNAs, with a Z-score above a given threshold. Our network approached scale freeness (i.e. approached the degree distribution of a random scale-free network generated by the Barabási-Albert model (Barabasi and Albert, 1999)) at a Z-score threshold of five, with higher thresholds not resulting in better approximations, and we therefore used this threshold for most of the analysis in this paper. Links based on MI values below 0.25 were deemed spurious and removed. We computed gene centrality for each gene in the network, including degree (number of neighbors), average neighborhood degree (the average degree of the neighbors), betweenness (the probability that the gene is part of the shortest network path between two arbitrary genes) and closeness (the shortest distance between a gene and all other genes in the network). The expression specificity score of a gene was calculated as the highest observed ratio between the average expression within and outside a zone of consecutive samples. All zones containing from three to 10 samples were considered and the final score was calculated as the average of the highest score from each replicate tree. The modular network was generated as explained in the figure text of Figure 3.

### AspWood

A web resource was built to allow the community easy access to the expression data (AspWood, http://aspwood.popgenie.org). The user can start with an existing list of genes, create a new list by text search or select one of the precomputed clusters. The tool will generate a page for the selected gene list with modules showing the expression profiles, the co-expression network, gene information, functional enrichments and the heatmap. The co-expression network is interactive and allows the user to explore the data further. The gene list can be exported and used in tools throughout the PlantGenIE platform (http://plantgenie.org, Sundell et al. (2015)). AspWood is built with HTML5 and JavaScript for user interfaces, and PhP and Python for more advanced server side functions.

## Acknowledgments

Thanks to Kjell Olofsson for performing the cryosectioning.

## Availability of data and materials

Raw RNA-Seq reads are available at the European Nucleotide Archive (ENA, http://www.ebi.ac.uk/ena/): accession number ERP016242. Processed data is available through the interactive AspWood web resource: http://aspwood.popgenie.org.

## Authors’ contributions

NRS, BS and TRH conceived and designed the experiment. MK prepared RNA for sequencing. DS implemented AspWood (with help from CM), and preformed the RNA-Seq preprocessing, the novel gene analysis (with help from ND) and coexpression network and enrichment analysis. TRH performed the cluster analysis. NRS, EJM, MK, CJ, VK, ON, HT, EP, UF, TN, BS and TRH analyzed and interpreted the data. All authors contributed text to the manuscript, and read and approved the final version.

## Additional figures and tables

### Additional figures

**Figure S1.**

Plots showing the expression diversity across the sampled series. Sample colors indicate the sample clusters from Figure 1B. (A) Box plots showing the expression distribution in each sample. The boxes indicate upper/lower quartiles (i.e. 25% of the genes are expressed higher/lower than the box) with the horizontal lines marking the median (i.e. 50% of the genes are expressed higher/lower than this value). The lines extending vertically from the boxes (whiskers) indicate the maximum/minimum values excluding outliers. Outliers (i.e. genes expressed more/less than 1.5 times the upper/lower quartile) are plotted as open circles. (B, C) The number of genes in each sample expressed above a threshold of 3 and 15, respectively. (D) The number of genes in each sample accounting for 90% of the expression in that sample.

**Figure S2.**

The sample clustering in Figure 1B with visible sample names (see Table S2).

**Figure S3.**

Marker genes. Expression is shown with (A) the variance stabilized transformation (VST) and (B) scaled counts per million (CPM, calculated as 2^VST^, scaled: mean centered and normalized by the standard deviation of each gene). PtSUS6/Potri.004G081300: *A. thaliana* sucrose synthase (*SUS*) genes *SUS5* and *SUS6* are known to be phloem localized (Barratt et al., 2009). The *Populus* homologs of *SUS5* and *SUS6* peaked in the two or three outermost sections of the single section series covering the cambial meristem, thus marking a phloem identity to these samples. The pooled phloem sample, consisting of six to seven section showed lower levels indicating that *SUS5* and *SUS6* expression decreased in the older phloem tissues. Expression was shown for one *SUS6* homolog (*PtSUS6*). PtCDC2/Potri.016G142800: The dividing meristem was marked by a set of cyclins typically known to be involved in the different stages of cell cycling. These markers indicated that sample cluster ***i*** and ***ii*** from the hierarchical clustering was split in the middle of the meristem. Expression was shown for *PtCDC2. PtEXPA1*/Potri.001G240900: Cell expansion was marked by the *Populus* alpha expansin (Gray-Mitsumune et al., 2004). The *PtEXPAl* expression marked the broader region covering both the phloem and xylem side, and decreased sharply at the border to sample cluster ***iii***. *PtCesA8-B*/Potri.004G059600: *CesA4, 7* and *8* are well-described hallmark genes for secondary wall formation in xylem cells (Kumar et al., 2009). The corresponding transcripts in aspen showed two distinct peaks, one minor on the phloem side marking secondary wall formation in phloem, and another major peak on the xylem side marking the bulk of secondary wall formation in xylem vessels, fibers and rays. Expression was shown for *PtCesA8-B. PtBFN1*/Potri.011G044500: Vessel cells undergo cell death before fiber cells. Xylem specific proteases and nucleases are responsible for the autolysis of xylem cells following cell death were selected to mark the timing of the cell death of fibers and vessel elements (reviewed by (Escamez and Tuominen, 2014)). In aspen, a homolog of the *A. thaliana* bifunctional nuclease 1 (BFN1) protein showed two peaks of expression; one appearing at the end of sample cluster ***iii*** and the other at the end of sample cluster ***iv***. This corresponds well with the locations of cell death, as estimated on the basis of the viability assays of the xylem tissues (TableS2). Thus, the sharp decrease in the expression of *PtBFN1* marks the border of living vessel elements and fibers, respectively.

**Figure S4.**

The hierarchical clustering from Figure 1BC, but with samples ordered according to sampling order for each of the four replicate trees and not according to expression similarity. The four sample clusters identified in Figure 1B are indicated by the color bar above the heatmap.

**Figure S5.**

Comparison of expression domains of proposed direct targets of the four *SND1* paralogs in wood-forming tissues (Potri.011G153300, Potri.001G448400, Potri.014G104800, Potri.002G178700). The heatmap shows expression profiles of *SND1s* (in green) as well as proposed direct targets of these identified in previous studies (in blue (Ye and Zhong, 2015) and black (Lin et al., 2013)). Genes proposed as targets by both previous studies are in red.

**Figure S6.**

Genotyping. RNA-Seq reads from the four AspWood replicates, as well as from four different genotypes from an independent experiment (controls), were used to call SNPs for all genes on Chromosome 1 of the *P. trichocarpa* genome. A Principle Component Analysis (PCA) showed that the four AspWood trees (turquoise) were clonal replicates and that the controls (red) were not.

### Additional tables

**Table S1.** Information about the sampled trees including height, diameter, age, and sampling height.

**Table S2.** Anatomical characterization of each section in the sample series, and description of pooling for RNA-Seq analysis. The colors indicate sample clusters according to Figure 1B. Cell viability measurements are also indicated (purple bars).

**Table S3.** Expression values for the 28,294 expressed, annotated genes (version 3 of the *P. trichocarpa* genome).

**Table S4.** Expression values for the 78 expressed, novel protein-coding genes identified in this study. Chromosomal positions in the *P. trichocarpa* genome is also indicated.

**Table S5.** Expression values for the 567 expressed, long non-coding RNAs identified in this study. Chromosomal positions in the *P. trichocarpa* genome is also indicated.

**Table S6.** Expression values for the 307 expressed, transcript fragments identified in this study. Chromosomal positions in the *P. trichocarpa* genome is also indicated.

**Table S7.** All 28,294 expressed, annotated protein-coding genes with cluster assignments according to the hierarchical clustering. Functional annotations are from www.phytozome.net.

**Table S8.** Network centrality statistics for the genes in the co-expression network at a co-expression threshold of five: 13,882 annotated genes (Sheet named *Annotated*), 25 novel coding genes (*Novel coding*), 205 novel long noncoding RNAs (*Novel non-coding*) and 87 transcript fragments (*Novel fragments*). Each sheet includes columns for network statistics, expression specificity (see Methods), Pearson correlations to the mean expression profiles of the gene clusters obtained by hierarchical clustering and annotations (www.phytozome.net). A separate sheet contains the same information for genes selected as representatives in the modular network (*Annotated representative*). Finally, there is a sheet providing statistical associations observed in the network (*Statistics*) and a sheet providing statistics on BLAST hit with E-value < 1E-10 to other species (*BLAST*).

**Table S9.** Central genes and enriched gene functions in the 41 network clusters.

**Table S10.** Complete lists of genes for the different biological stories.

**Table S11.** Analysis of WGD paralogs. (A) Expression correlation for all paralog pairs, where at least one member of the pair was expressed in the section series. (B) Numbers of pairs (with p-values) shared between the clusters. Table with the number of diverged pairs after applying different correlation thresholds.

**Table S12.** RNA-seq data quality. (A) Read counts for each sample including the number of raw reads and the number of reads after filtering, trimming and alignment (in millions). (B) Principle Component Analysis (PCA) of all samples based on the variance stabilized expression data.

## References

Allwright, M.R., Payne, A., Emiliani, G., Milner, S., Viger, M., Rouse, F., Keurentjes, J.J., Berard, A., Wildhagen, H., Faivre-Rampant, P., Polle, A., Morgante, M., and Taylor, G. (2016). Biomass traits and candidate genes for bioenergy revealed through association genetics in coppiced European Populus nigra (L.). Biotechnol Biofuels 9, 195.

Anders, S., Pyl, P.T., and Huber, W. (2014). HTSeq; A Python framework to work with high-throughput sequencing data.

Aspeborg, H., Schrader, J., Coutinho, P.M., Stam, M., Kallas, A., Djerbi, S., Nilsson, P., Denman, S., Amini, B., Sterky, F., Master, E., Sandberg, G., Mellerowicz, E., Sundberg, B., Henrissat, B., and Teeri, T.T. (2005). Carbohydrate-active enzymes involved in the secondary cell wall biogenesis in hybrid aspen. Plant Physiol 137, 983-997.

Atmodjo, M.A., Sakuragi, Y., Zhu, X., Burrell, A.J., Mohanty, S.S., Atwood, J.A., 3rd, Orlando, R., Scheller, H.V., and Mohnen, D. (2011). Galacturonosyltransferase (GAUT)1 and GAUT7 are the core of a plant cell wall pectin biosynthetic homogalacturonan:galacturonosyltransferase complex. Proc Natl Acad Sci U S A 108, 20225-20230.

Barabasi, A.L., and Albert, R. (1999). Emergence of scaling in random networks. Science 286, 509-512.

Barnett, J.R. (1981). Secondary xylem cell development. In Xylem Cell Development, J.R. Barnett, ed (London, England: Castle House), pp. 47-95.

Barratt, D.H.P., Derbyshire, P., Findlay, K., Pike, M., Wellner, N., Lunn, J., Feil, R., Simpson, C., Maule, A.J., and Smith, A.M. (2009). Normal growth of Arabidopsis requires cytosolic invertase but not sucrose synthase. P Natl Acad Sci USA 106, 13124-13129.

Barros, J., Serk, H., Granlund, I., and Pesquet, E. (2015). The cell biology of lignification in higher plants. Ann Bot 115, 1053-1074.

Biswal, A.K., Soeno, K., Gandla, M.L., Immerzeel, P., Pattathil, S., Lucenius, J., Serimaa, R., Hahn, M.G., Moritz, T., Jonsson, L.J., Israelsson-Nordstrom, M., and Mellerowicz, E.J. (2014). Aspen pectate lyase PtxtPL1-27 mobilizes matrix polysaccharides from woody tissues and improves saccharification yield. Biotechnol Biofuels 7, 11.

Boerjan, W., Ralph, J., and Baucher, M. (2003). Lignin biosynthesis. Annu Rev Plant Biol 54, 519-546.

Bollhöner, B., Prestele, J., and Tuominen, H. (2012). Xylem cell death: emerging understanding of regulation and function. J Exp Bot 63, 1081-1094.

Bollhöner, B., Zhang, B., Stael, S., Denance, N., Overmyer, K., Goffner, D., Van Breusegem, F., and Tuominen, H. (2013). Post mortem function of AtMC9 in xylem vessel elements. New Phytol 200, 498-510.

Bonke, M., Thitamadee, S., Mahonen, A.P., Hauser, M.T., and Helariutta, Y. (2003). APL regulates vascular tissue identity in Arabidopsis. Nature 426, 181-186.

Bourquin, V., Nishikubo, N., Abe, H., Brumer, H., Denman, S., Eklund, M., Christiernin, M., Teeri, T.T., Sundberg, B., and Mellerowicz, E.J. (2002). Xyloglucan endotransglycosylases have a function during the formation of secondary cell walls of vascular tissues. Plant Cell 14, 3073-3088.

Carroll, A., Mansoori, N., Li, S., Lei, L., Vernhettes, S., Visser, R.G., Somerville, C., Gu, Y., and Trindade, L.M. (2012). Complexes with mixed primary and secondary cellulose synthases are functional in Arabidopsis plants. Plant Physiol 160, 726-737.

Cavalier, D.M., and Keegstra, K. (2006). Two xyloglucan xylosyltransferases catalyze the addition of multiple xylosyl residues to cellohexaose. J Biol Chem 281, 34197-34207.

Chou, Y.H., Pogorelko, G., and Zabotina, O.A. (2012). Xyloglucan xylosyltransferases XXT1, XXT2, and XXT5 and the glucan synthase CSLC4 form Golgi-localized multiprotein complexes. Plant Physiol 159, 1355-1366.

Christensen, J.H., Bauw, G., Welinder, K.G., Van Montagu, M., and Boerjan, W. (1998). Purification and characterization of peroxidases correlated with lignification in poplar xylem. Plant Physiol 118, 125-135.

Courtois-Moreau, C.L., Pesquet, E., Sjodin, A., Muniz, L., Bollhöner, B., Kaneda, M., Samuels, L., Jansson, S., and Tuominen, H. (2009). A unique program for cell death in xylem fibers of Populus stem. Plant J 58, 260-274.

Dai, X., Hu, Q., Cai, Q., Feng, K., Ye, N., Tuskan, G.A., Milne, R., Chen, Y., Wan, Z., Wang, Z., Luo, W., Wang, K., Wan, D., Wang, M., Wang, J., Liu, J., and Yin, T. (2014). The willow genome and divergent evolution from poplar after the common genome duplication. Cell Res 24, 1274-1277.

De Rybel, B., Moller, B., Yoshida, S., Grabowicz, I., Barbier de Reuille, P., Boeren, S., Smith, R.S., Borst, J.W., and Weijers, D. (2013). A bHLH complex controls embryonic vascular tissue establishment and indeterminate growth in Arabidopsis. Dev Cell 24, 426-437.

De Rybel, B., Adibi, M., Breda, A.S., Wendrich, J.R., Smit, M.E., Novak, O., Yamaguchi, N., Yoshida, S., Van Isterdael, G., Palovaara, J., Nijsse, B., Boekschoten, M.V., Hooiveld, G., Beeckman, T., Wagner, D., Ljung, K., Fleck, C., and Weijers, D. (2014). Integration of growth and patterning during vascular tissue formation in Arabidopsis. Science 345, 1255215.

Delhomme, N., Mähler, N., Schiffthaler, B., Sundell, D., Mannapperuma, C., Hvidsten, T., and Street, N. (2014). Guidelines for RNA-Seq data analysis. EpiGeneSys Protocol.

Depuydt, S., Rodriguez-Villalon, A., Santuari, L., Wyser-Rmili, C., Ragni, L., and Hardtke, C.S. (2013). Suppression of Arabidopsis protophloem differentiation and root meristem growth by CLE45 requires the receptor-like kinase BAM3. Proc Natl Acad Sci U S A 110, 7074-7079.

Derbyshire, P., Menard, D., Green, P., Saalbach, G., Buschmann, H., Lloyd, C.W., and Pesquet, E. (2015). Proteomic Analysis of Microtubule Interacting Proteins over the Course of Xylem Tracheary Element Formation in Arabidopsis. Plant Cell 27, 2709-2726.

Dobin, A., Davis, C.A., Schlesinger, F., Drenkow, J., Zaleski, C., Jha, S., Batut, P., Chaisson, M., and Gingeras, T.R. (2013). STAR: ultrafast universal RNA-seq aligner. Bioinformatics 29, 15-21.

Egelund, J., Petersen, B.L., Motawia, M.S., Damager, I., Faik, A., Olsen, C.E., Ishii, T., Clausen, H., Ulvskov, P., and Geshi, N. (2006). Arabidopsis thaliana RGXT1 and RGXT2 encode Golgi-localized (1,3)-alpha-D-xylosyltransferases involved in the synthesis of pectic rhamnogalacturonan-II. Plant Cell 18, 2593-2607.

Emery, J.F., Floyd, S.K., Alvarez, J., Eshed, Y., Hawker, N.P., Izhaki, A., Baum, S.F., and Bowman, J.L. (2003). Radial patterning of Arabidopsis shoots by class III HD-ZIP and KANADI genes. Curr Biol 13, 1768-1774.

Endo, S., Pesquet, E., Yamaguchi, M., Tashiro, G., Sato, M., Toyooka, K., Nishikubo, N., Udagawa-Motose, M., Kubo, M., Fukuda, H., and Demura, T. (2009). Identifying new components participating in the secondary cell wall formation of vessel elements in zinnia and Arabidopsis. Plant Cell 21, 11551165.

Escamez, S., and Tuominen, H. (2014). Programmes of cell death and autolysis in tracheary elements: when a suicidal cell arranges its own corpse removal. J Exp Bot 65, 1313-1321.

Espinosa-Ruiz, A., Saxena, S., Schmidt, J., Mellerowicz, E., Miskolczi, P., Bako, L., and Bhalerao, R.P. (2004). Differential stage-specific regulation of cyclin-dependent kinases during cambial dormancy in hybrid aspen. Plant J 38, 603-615.

Etchells, J.P., and Turner, S.R. (2010). The PXY-CLE41 receptor ligand pair defines a multifunctional pathway that controls the rate and orientation of vascular cell division. Development 137, 767-774.

Evert, R.F., and Eichhorn, S.E. (2006). In Esau’s Plant Anatomy (Willey & Sons, Inc), pp. 407-425.

Faith, J.J., Hayete, B., Thaden, J.T., Mogno, I., Wierzbowski, J., Cottarel, G., Kasif, S., Collins, J.J., and Gardner, T.S. (2007). Large-scale mapping and validation of Escherichia coli transcriptional regulation from a compendium of expression profiles. PLoS biology 5, e8.

Fujii, T., Harada, H., and Saiki, H. (1981). Ultrastructure of ‘amorphous layer’ in xylem parenchyma cell wall of angiosperm species. Mokuzai Gakkaishi 27, 149–156.

Fukuda, H. (2016). Signaling, transcriptional regulation, and asynchronous pattern formation governing plant xylem development. Proc Jpn Acad Ser B Phys Biol Sci 92, 98-107.

Furuta, K.M., Yadav, S.R., Lehesranta, S., Belevich, I., Miyashima, S., Heo, J.O., Vaten, A., Lindgren, O., De Rybel, B., Van Isterdael, G., Somervuo, P., Lichtenberger, R., Rocha, R., Thitamadee, S., Tahtiharju, S., Auvinen, P., Beeckman, T., Jokitalo, E., and Helariutta, Y. (2014). Arabidopsis NAC45/86 direct sieve element morphogenesis culminating in enucleation. Science 345, 933-937.

Goodstein, D.M., Shu, S., Howson, R., Neupane, R., Hayes, R.D., Fazo, J., Mitros, T., Dirks, W., Hellsten, U., Putnam, N., and Rokhsar, D.S. (2012). Phytozome: a comparative platform for green plant genomics. Nucleic Acids Res 40, D1178-1186.

Goodwin, S., McPherson, J.D., and McCombie, W.R. (2016). Coming of age: ten years of next-generation sequencing technologies. Nat Rev Genet 17, 333-351.

Gouzy, J., Carrere, S., and Schiex, T. (2009). FrameDP: sensitive peptide detection on noisy matured sequences. Bioinformatics 25, 670-671.

Grabherr, M.G., Haas, B.J., Yassour, M., Levin, J.Z., Thompson, D.A., Amit, I., Adiconis, X., Fan, L., Raychowdhury, R., Zeng, Q., Chen, Z., Mauceli, E., Hacohen, N., Gnirke, A., Rhind, N., di Palma, F., Birren, B.W., Nusbaum, C., Lindblad-Toh, K., Friedman, N., and Regev, A. (2011). Full-length transcriptome assembly from RNA-Seq data without a reference genome. Nat Biotechnol 29, 644-652.

Gray-Mitsumune, M., Blomquist, K., McQueen-Mason, S., Teeri, T.T., Sundberg, B., and Mellerowicz, E.J. (2008). Ectopic expression of a wood-abundant expansin PttEXPA1 promotes cell expansion in primary and secondary tissues in aspen. Plant Biotechnol J 6, 62-72.

Gray-Mitsumune, M., Mellerowicz, E.J., Abe, H., Schrader, J., Winzell, A., Sterky, F., Blomqvist, K., McQueen-Mason, S., Teeri, T.T., and Sundberg, B. (2004). Expansins abundant in secondary xylem belong to subgroup A of the alpha-expansin gene family. Plant Physiol 135, 1552-1564.

Haas, B.J., Delcher, A.L., Mount, S.M., Wortman, J.R., Smith, R.K., Jr., Hannick, L.I., Maiti, R., Ronning, C.M., Rusch, D.B., Town, C.D., Salzberg, S.L., and White, O. (2003). Improving the Arabidopsis genome annotation using maximal transcript alignment assemblies. Nucleic Acids Res 31, 5654-5666.

Harholt, J., Jensen, J.K., Sorensen, S.O., Orfila, C., Pauly, M., and Scheller, H.V. (2006). ARABINAN DEFICIENT 1 is a putative arabinosyltransferase involved in biosynthesis of pectic arabinan in Arabidopsis. Plant Physiol 140, 49-58.

Hermansen, R.A., Hvidsten, T.R., Sandve, S.R., and Liberles, D.A. (2016). Extracting functional trends from whole genome duplication events using comparative genomics. Biol Proced Online 18, 11.

Herrero, J., Fernandez-Perez, F., Yebra, T., Novo-Uzal, E., Pomar, F., Pedreno, M.A., Cuello, J., Guera, A., Esteban-Carrasco, A., and Zapata, J.M. (2013). Bioinformatic and functional characterization of the basic peroxidase 72 from Arabidopsis thaliana involved in lignin biosynthesis. Planta 237, 1599-1612.

Hertzberg, M., Aspeborg, H., Schrader, J., Andersson, A., Erlandsson, R., Blomqvist, K., Bhalerao, R., Uhlen, M., Teeri, T.T., Lundeberg, J., Sundberg, B., Nilsson, P., and Sandberg, G. (2001). A transcriptional roadmap to wood formation. Proc Natl Acad Sci U S A 98, 14732-14737.

Hill, J.L., Jr., Hammudi, M.B., and Tien, M. (2014). The Arabidopsis cellulose synthase complex: a proposed hexamer of CESA trimers in an equimolar stoichiometry. Plant Cell 26, 4834-4842.

Hirakawa, Y., Shinohara, H., Kondo, Y., Inoue, A., Nakanomyo, I., Ogawa, M., Sawa, S., Ohashi-Ito, K., Matsubayashi, Y., and Fukuda, H. (2008). Noncell-autonomous control of vascular stem cell fate by a CLE peptide/receptor system. Proc Natl Acad Sci U S A 105, 15208-15213.

Ilegems, M., Douet, V., Meylan-Bettex, M., Uyttewaal, M., Brand, L., Bowman, J.L., and Stieger, P.A. (2010). Interplay of auxin, KANADI and Class III HD-ZIP transcription factors in vascular tissue formation. Development 137, 975-984.

Immanen, J., Nieminen, K., Duchens Silva, H., Rodriguez Rojas, F., Meisel, L.A., Silva, H., Albert, V.A., Hvidsten, T.R., and Helariutta, Y. (2013). Characterization of cytokinin signaling and homeostasis gene families in two hardwood tree species: Populus trichocarpa and Prunus persica. BMC Genomics 14, 885.

Immanen, J., Nieminen, K., Smolander, O.P., Kojima, M., Alonso Serra, J., Koskinen, P., Zhang, J., Elo, A., Mahonen, A.P., Street, N., Bhalerao, R.P., Paulin, L., Auvinen, P., Sakakibara, H., and Helariutta, Y. (2016). Cytokinin and Auxin Display Distinct but Interconnected Distribution and Signaling Profiles to Stimulate Cambial Activity. Curr Biol 26, 1990-1997.

Ito, J., and Fukuda, H. (2002). ZEN1 is a key enzyme in the degradation of nuclear DNA during programmed cell death of tracheary elements. Plant Cell 14, 3201-3211.

Jensen, J.K., Johnson, N.R., and Wilkerson, C.G. (2014). Arabidopsis thaliana IRX10 and two related proteins from psyllium and Physcomitrella patens are xylan xylosyltransferases. Plant J 80, 207-215.

Johnsson, C., and Fischer, U. (2016). Cambial stem cells and their niche. Plant Sci 252, 239-245.

Kang, I.H., Steffen, J.G., Portereiko, M.F., Lloyd, A., and Drews, G.N. (2008). The AGL62 MADS domain protein regulates cellularization during endosperm development in Arabidopsis. Plant Cell 20, 635-647.

Kubo, M., Udagawa, M., Nishikubo, N., Horiguchi, G., Yamaguchi, M., Ito, J., Mimura, T., Fukuda, H., and Demura, T. (2005). Transcription switches for protoxylem and metaxylem vessel formation. Genes Dev 19, 1855-1860.

Kumar, M., Thammannagowda, S., Bulone, V., Chiang, V., Han, K.H., Joshi, C.P., Mansfield, S.D., Mellerowicz, E., Sundberg, B., Teeri, T., and Ellis, B.E. (2009). An update on the nomenclature for the cellulose synthase genes in Populus. Trends Plant Sci 14, 248-254.

Larson, P.R. (1994). The Vascular Cambium: Development and Structure. (Berlin: Springer-Verlag).

Lee, C., Teng, Q., Zhong, R., and Ye, Z.H. (2011). Molecular dissection of xylan biosynthesis during wood formation in poplar. Mol Plant 4, 730-747.

Li, G., Gelernter, J., Kranzler, H.R., and Zhao, H. (2012). M(3): an improved SNP calling algorithm for Illumina BeadArray data. Bioinformatics 28, 358-365.

Li, Y., Kajita, S., Kawai, S., Katayama, Y., and Morohoshi, N. (2003). Down-regulation of an anionic peroxidase in transgenic aspen and its effect on lignin characteristics. J Plant Res 116, 175-182.

Lin, Y.C., Li, W., Sun, Y.H., Kumari, S., Wei, H., Li, Q., Tunlaya-Anukit, S., Sederoff, R.R., and Chiang, V.L. (2013). SND1 transcription factor-directed quantitative functional hierarchical genetic regulatory network in wood formation in Populus trichocarpa. Plant Cell 25, 4324-4341.

Liwanag, A.J., Ebert, B., Verhertbruggen, Y., Rennie, E.A., Rautengarten, C., Oikawa, A., Andersen, M.C., Clausen, M.H., and Scheller, H.V. (2012). Pectin biosynthesis: GALS1 in Arabidopsis thaliana is a beta-1,4-galactan beta-1,4-galactosyltransferase. Plant Cell 24, 5024-5036.

Love, M.I., Huber, W., and Anders, S. (2014). Moderated estimation of fold change and dispersion for RNA-seq data with DESeq2. Genome Biol 15, 550.

Lu, S., Li, Q., Wei, H., Chang, M.J., Tunlaya-Anukit, S., Kim, H., Liu, J., Song, J., Sun, Y.H., Yuan, L., Yeh, T.F., Peszlen, I., Ralph, J., Sederoff, R.R., and Chiang, V.L. (2013). Ptr-miR397a is a negative regulator of laccase genes affecting lignin content in Populus trichocarpa. Proc Natl Acad Sci U S A 110, 10848-10853.

McCarthy, R.L., Zhong, R., Fowler, S., Lyskowski, D., Piyasena, H., Carleton, K., Spicer, C., and Ye, Z.H. (2010). The poplar MYB transcription factors, PtrMYB3 and PtrMYB20, are involved in the regulation of secondary wall biosynthesis. Plant Cell Physiol 51, 1084-1090.

Mellerowicz, E.J., and Gorshkova, T.A. (2012). Tensional stress generation in gelatinous fibres: a review and possible mechanism based on cell-wall structure and composition. J Exp Bot 63, 551-565.

Mellerowicz, E.J., Baucher, M., Sundberg, B., and Boerjan, W. (2001). Unravelling cell wall formation in the woody dicot stem. Plant Mol Biol 47, 239-274.

Moreau, C., Aksenov, N., Lorenzo, M.G., Segerman, B., Funk, C., Nilsson, P., Jansson, S., and Tuominen, H. (2005). A genomic approach to investigate developmental cell death in woody tissues of Populus trees. Genome Biol 6, R34.

Mortimer, J.C., Faria-Blanc, N., Yu, X., Tryfona, T., Sorieul, M., Ng, Y.Z., Zhang, Z., Stott, K., Anders, N., and Dupree, P. (2015). An unusual xylan in Arabidopsis primary cell walls is synthesised by GUX3, IRX9L, IRX10L and IRX14. Plant J 83, 413-426.

Murakami, R., Funada, R., Sano, Y., and Ohtani, J. (1999). The Differentiation of Contact Cells and Isolation Cells in the Xylem Ray Parenchyma of Populus maximowiczii. Annals of Botany 84, 429-435.

Mutwil, M., Klie, S., Tohge, T., Giorgi, F.M., Wilkins, O., Campbell, M.M., Fernie, A.R., Usadel, B., Nikoloski, Z., and Persson, S. (2011). PlaNet: combined sequence and expression comparisons across plant networks derived from seven species. Plant Cell 23, 895-910.

Nakaba, S., Begum, S., Yamagishi, Y., Jin, H.-O., Kubo, T., and Funada, R. (2012). Differences in the timing of cell death, differentiation and function among three different types of ray parenchyma cells in the hardwood Populus sieboldii × P. grandidentata. Trees 26, 743.

Netotea, S., Sundell, D., Street, N.R., and Hvidsten, T.R. (2014). ComPlEx: conservation and divergence of co-expression networks in A. thaliana, Populus and O. sativa. BMC Genomics 15, 106.

Ohashi-Ito, K., Saegusa, M., Iwamoto, K., Oda, Y., Katayama, H., Kojima, M., Sakakibara, H., and Fukuda, H. (2014). A bHLH complex activates vascular cell division via cytokinin action in root apical meristem. Curr Biol 24, 2053-2058.

Oikawa, A., Lund, C.H., Sakuragi, Y., and Scheller, H.V. (2013). Golgi-localized enzyme complexes for plant cell wall biosynthesis. Trends Plant Sci 18, 49-58.

Proost, S., Fostier, J., De Witte, D., Dhoedt, B., Demeester, P., Van de Peer, Y., and Vandepoele, K. (2012). i-ADHoRe 3.0--fast and sensitive detection of genomic homology in extremely large data sets. Nucleic Acids Res 40, e11.

Quinlan, A.R., and Hall, I.M. (2010). BEDTools: a flexible suite of utilities for comparing genomic features. Bioinformatics 26, 841-842.

R-Core-Team. (2012). R: A Language and Environment for Statistical Computing (Vienna, 780 Austria OR - R Foundation for Statistical Computing).

Ranocha, P., McDougall, G., Hawkins, S., Sterjiades, R., Borderies, G., Stewart, D., Cabanes-Macheteau, M., Boudet, A.M., and Goffner, D. (1999). Biochemical characterization, molecular cloning and expression of laccases - a divergent gene family - in poplar. Eur J Biochem 259, 485-495.

Ratke, C., Pawar, P.M., Balasubramanian, V.K., Naumann, M., Duncranz, M.L., Derba-Maceluch, M., Gorzsas, A., Endo, S., Ezcurra, I., and Mellerowicz, E. J. (2015). Populus GT43 family members group into distinct sets required for primary and secondary wall xylan biosynthesis and include useful promoters for wood modification. Plant Biotechnol J 13, 26-37.

Rodgers-Melnick, E., Mane, S.P., Dharmawardhana, P., Slavov, G.T., Crasta, O.R., Strauss, S.H., Brunner, A.M., and Difazio, S.P. (2012). Contrasting patterns of evolution following whole genome versus tandem duplication events in Populus. Genome Res 22, 95-105.

Rodriguez-Villalon, A., Gujas, B., Kang, Y.H., Breda, A.S., Cattaneo, P., Depuydt, S., and Hardtke, C.S. (2014). Molecular genetic framework for protophloem formation. Proc Natl Acad Sci U S A 111, 11551-11556.

Rosin, F.M., and Kramer, E.M. (2009). Old dogs, new tricks: regulatory evolution in conserved genetic modules leads to novel morphologies in plants. Dev Biol 332, 25-35.

Ruzicka, K., Ursache, R., Hejatko, J., and Helariutta, Y. (2015). Xylem development - from the cradle to the grave. New Phytol 207, 519-535.

Sasaki, S., Nishida, T., Tsutsumi, Y., and Kondo, R. (2004). Lignin dehydrogenative polymerization mechanism: a poplar cell wall peroxidase directly oxidizes polymer lignin and produces in vitro dehydrogenative polymer rich in beta-O-4 linkage. FEBS Lett 562, 197-201.

Sasaki, S., Nonaka, D., Wariishi, H., Tsutsumi, Y., and Kondo, R. (2008). Role of Tyr residues on the protein surface of cationic cell-wall-peroxidase (CWPO-C) from poplar: potential oxidation sites for oxidative polymerization of lignin. Phytochemistry 69, 348-355.

Scacchi, E., Salinas, P., Gujas, B., Santuari, L., Krogan, N., Ragni, L., Berleth, T., and Hardtke, C.S. (2010). Spatio-temporal sequence of cross-regulatory events in root meristem growth. Proc Natl Acad Sci U S A 107, 22734-22739.

Schlereth, A., Moller, B., Liu, W., Kientz, M., Flipse, J., Rademacher, E.H., Schmid, M., Jurgens, G., and Weijers, D. (2010). MONOPTEROS controls embryonic root initiation by regulating a mobile transcription factor. Nature 464, 913-916.

Schrader, J., Nilsson, J., Mellerowicz, E., Berglund, A., Nilsson, P., Hertzberg, M., and Sandberg, G. (2004). A high-resolution transcript profile across the wood-forming meristem of poplar identifies potential regulators of cambial stem cell identity. Plant Cell 16, 2278-2292.

Shi, R., Sun, Y.H., Li, Q., Heber, S., Sederoff, R., and Chiang, V.L. (2010). Towards a systems approach for lignin biosynthesis in Populus trichocarpa: transcript abundance and specificity of the monolignol biosynthetic genes. Plant Cell Physiol 51, 144-163.

Shi, R., Wang, J.P., Lin, Y.C., Li, Q., Sun, Y.H., Chen, H., Sederoff, R.R., and Chiang, V.L. (2017). Tissue and cell-type co-expression networks of transcription factors and wood component genes in Populus trichocarpa. Planta.

Shigeto, J., Itoh, Y., Hirao, S., Ohira, K., Fujita, K., and Tsutsumi, Y. (2015). Simultaneously disrupting AtPrx2, AtPrx25 and AtPrx71 alters lignin content and structure in Arabidopsis stem. J Integr Plant Biol 57, 349-356.

Song, D., Shen, J., and Li, L. (2010). Characterization of cellulose synthase complexes in Populus xylem differentiation. New Phytol 187, 777-790.

Sterck, L., Rombauts, S., Jansson, S., Sterky, F., Rouze, P., and Van de Peer, Y. (2005). EST data suggest that poplar is an ancient polyploid. New Phytol 167, 165-170.

Sundell, D., Mannapperuma, C., Netotea, S., Delhomme, N., Lin, Y.C., Sjodin, A., Van de Peer, Y., Jansson, S., Hvidsten, T.R., and Street, N.R. (2015). The Plant Genome Integrative Explorer Resource: PlantGenIE.org. New Phytol 208, 1149-1156.

Takata, N., and Taniguchi, T. (2015). Expression divergence of cellulose synthase (CesA) genes after a recent whole genome duplication event in Populus. Planta 241, 29-42.

Trapnell, C., Williams, B.A., Pertea, G., Mortazavi, A., Kwan, G., van Baren, M.J., Salzberg, S.L., Wold, B.J., and Pachter, L. (2010). Transcript assembly and quantification by RNA-Seq reveals unannotated transcripts and isoform switching during cell differentiation. Nat Biotechnol 28, 511-515.

Truernit, E., Bauby, H., Belcram, K., Barthelemy, J., and Palauqui, J.C. (2012). OCTOPUS, a polarly localised membrane-associated protein, regulates phloem differentiation entry in Arabidopsis thaliana. Development 139, 1306-1315.

Tuskan, G.A., DiFazio, S., Jansson, S., Bohlmann, J., Grigoriev, I., Hellsten, U., Putnam, N., Ralph, S., Rombauts, S., Salamov, A., Schein, J., Sterck, L., Aerts, A., Bhalerao, R.R., Bhalerao, R.P., Blaudez, D., Boerjan, W., Brun, A., Brunner, A., Busov, V., Campbell, M., Carlson, J., Chalot, M., Chapman, J., Chen, G.L., Cooper, D., Coutinho, P.M., Couturier, J., Covert, S., Cronk, Q., Cunningham, R., Davis, J., Degroeve, S., Dejardin, A., Depamphilis, C., Detter, J., Dirks, B., Dubchak, I., Duplessis, S., Ehlting, J., Ellis, B., Gendler, K., Goodstein, D., Gribskov, M., Grimwood, J., Groover, A., Gunter, L., Hamberger, B., Heinze, B., Helariutta, Y., Henrissat, B., Holligan, D., Holt, R., Huang, W., Islam-Faridi, N., Jones, S., Jones-Rhoades, M., Jorgensen, R., Joshi, C., Kangasjarvi, J., Karlsson, J., Kelleher, C., Kirkpatrick, R., Kirst, M., Kohler, A., Kalluri, U., Larimer, F., Leebens-Mack, J., Leple, J.C., Locascio, P., Lou, Y., Lucas, S., Martin, F., Montanini, B., Napoli, C., Nelson, D.R., Nelson, C., Nieminen, K., Nilsson, O., Pereda, V., Peter, G., Philippe, R., Pilate, G., Poliakov, A., Razumovskaya, J., Richardson, P., Rinaldi, C., Ritland, K., Rouze, P., Ryaboy, D., Schmutz, J., Schrader, J., Segerman, B., Shin, H., Siddiqui, A., Sterky, F., Terry, A., Tsai, C.J., Uberbacher, E., Unneberg, P., Vahala, J., Wall, K., Wessler, S., Yang, G., Yin, T., Douglas, C., Marra, M., Sandberg, G., Van de Peer, Y., and Rokhsar, D. (2006). The genome of black cottonwood, Populus trichocarpa (Torr. & Gray). Science 313, 1596-1604.

Uggla, C., and Sundberg, B. (2002). Chapter 13. Sampling of Cambial Region Tissues for High Resolution Analysis. In Wood Formation in Trees (CRC Press), pp. 215–228.

Uggla, C., Moritz, T., Sandberg, G., and Sundberg, B. (1996). Auxin as a positional signal in pattern formation in plants. Proc Natl Acad Sci U S A 93, 9282-9286.

Ulitsky, I. (2016). Evolution to the rescue: using comparative genomics to understand long non-coding RNAs. Nat Rev Genet 17, 601-614.

Urbanowicz, B.R., Pena, M.J., Moniz, H.A., Moremen, K.W., and York, W.S. (2014). Two Arabidopsis proteins synthesize acetylated xylan in vitro. Plant J 80, 197-206.

Wang, H.H., Tang, R.J., Liu, H., Chen, H.Y., Liu, J.Y., Jiang, X.N., and Zhang, H.X. (2013). Chimeric repressor of PtSND2 severely affects wood formation in transgenic Populus. Tree physiology 33, 878-886.

Wullschleger, S.D., Weston, D.J., DiFazio, S.P., and Tuskan, G.A. (2013). Revisiting the sequencing of the first tree genome: Populus trichocarpa. Tree physiology 33, 357-364.

Yamaguchi, M., Kubo, M., Fukuda, H., and Demura, T. (2008). Vascular-related NAC-DOMAIN7 is involved in the differentiation of all types of xylem vessels in Arabidopsis roots and shoots. Plant J 55, 652-664.

Ye, Z.H., and Zhong, R. (2015). Molecular control of wood formation in trees. J Exp Bot 66, 4119-4131.

Yoo, M.J., Liu, X., Pires, J.C., Soltis, P.S., and Soltis, D.E. (2014). Nonadditive gene expression in polyploids. Annu Rev Genet 48, 485-517.

Zabotina, O.A., van de Ven, W.T., Freshour, G., Drakakaki, G., Cavalier, D., Mouille, G., Hahn, M.G., Keegstra, K., and Raikhel, N.V. (2008). Arabidopsis XXT5 gene encodes a putative alpha-1,6-xylosyltransferase that is involved in xyloglucan biosynthesis. Plant J 56, 101-115.

Zeng, W., Lampugnani, E.R., Picard, K.L., Song, L., Wu, A.M., Farion, I.M., Zhao, J., Ford, K., Doblin, M.S., and Bacic, A. (2016). Asparagus IRX9, IRX10, and IRX14A Are Components of an Active Xylan Backbone Synthase Complex that Forms in the Golgi Apparatus. Plant Physiol 171, 93-109.

Zhang, W., Landback, P., Gschwend, A.R., Shen, B., and Long, M. (2015). New genes drive the evolution of gene interaction networks in the human and mouse genomes. Genome Biol 16, 202.

Zhao, Q., Nakashima, J., Chen, F., Yin, Y., Fu, C., Yun, J., Shao, H., Wang, X., Wang, Z.Y., and Dixon, R.A. (2013). Laccase is necessary and nonredundant with peroxidase for lignin polymerization during vascular development in Arabidopsis. Plant Cell 25, 3976-3987.

Zheng, X., Levine, D., Shen, J., Gogarten, S.M., Laurie, C., and Weir, B.S. (2012). A high-performance computing toolset for relatedness and principal component analysis of SNP data. Bioinformatics 28, 3326-3328.

Zhong, R., Demura, T., and Ye, Z.H. (2006). SND1, a NAC domain transcription factor, is a key regulator of secondary wall synthesis in fibers of Arabidopsis. Plant Cell 18, 3158-3170.

Zhong, R., McCarthy, R.L., Haghighat, M., and Ye, Z.H. (2013). The poplar MYB master switches bind to the SMRE site and activate the secondary wall biosynthetic program during wood formation. PLoS One 8, e69219.

Zhou, J., Zhong, R., and Ye, Z.H. (2014). Arabidopsis NAC domain proteins, VND1 to VND5, are transcriptional regulators of secondary wall biosynthesis in vessels. PLoS One 9, e105726.

